# LinearTurboFold: Linear-Time Global Prediction of Conserved Structures for RNA Homologs with Applications to SARS-CoV-2

**DOI:** 10.1101/2020.11.23.393488

**Authors:** Sizhen Li, He Zhang, Liang Zhang, Kaibo Liu, Boxiang Liu, David H. Mathews, Liang Huang

**Author notes:** Author contributions: L.H. and D.H.M. conceived the idea and directed the project. S.L., H.Z., L.H., and D.H.M. designed the algorithm; S.L. implemented it. D.H.M. guided the evaluation that S.L. and L.Z. carried out. S.L. and H.Z. wrote the manuscript; L.H., and D.H.M. revised it. L.K. made the webserver. H.Z., B.L., L.H., and D.H.M. guided the analysis on SARS-CoV-2.

## Abstract

The constant emergence of COVID-19 variants reduces the effectiveness of existing vaccines and test kits. Therefore, it is critical to identify conserved structures in SARS-CoV-2 genomes as potential targets for variant-proof diagnostics and therapeutics. However, the algorithms to predict these conserved structures, which simultaneously fold and align multiple RNA homologs, scale at best cubically with sequence length, and are thus infeasible for coronaviruses, which possess the longest genomes (∼30,000 *nt*) among RNA viruses. As a result, existing efforts on modeling SARS-CoV-2 structures resort to single sequence folding as well as local folding methods with short window sizes, which inevitably neglect long-range interactions that are crucial in RNA functions. Here we present LinearTurboFold, an efficient algorithm for folding RNA homologs that scales *linearly* with sequence length, enabling unprecedented *global* structural analysis on SARS-CoV-2. Surprisingly, on a group of SARS-CoV-2 and SARS-related genomes, LinearTurboFold’s purely *in silico* prediction not only is close to experimentally-guided models for local structures, but also goes far beyond them by capturing the end-to-end pairs between 5’ and 3’ UTRs (∼29,800 *nt* apart) that match perfectly with a purely experimental work. Furthermore, LinearTurboFold identifies novel conserved structures and conserved accessible regions as potential targets for designing efficient and mutation-insensitive small-molecule drugs, antisense oligonucleotides, siRNAs, CRISPR-Cas13 guide RNAs and RT-PCR primers. LinearTurboFold is a general technique that can also be applied to other RNA viruses and full-length genome studies, and will be a useful tool in fighting the current and future pandemics.

**Significance Statement:** Conserved RNA structures are critical for designing diagnostic and therapeutic tools for many diseases including COVID-19. However, existing algorithms are much too slow to model the global structures of full-length RNA viral genomes. We present LinearTurboFold, a linear-time algorithm that is orders of magnitude faster, making it the first method to simultaneously fold and align whole genomes of SARS-CoV-2 variants, the longest known RNA virus (∼30 kilobases). Our work enables unprecedented global structural analysis and captures long-range interactions that are out of reach for existing algorithms but crucial for RNA functions. LinearTurboFold is a general technique for full-length genome studies and can help fight the current and future pandemics.

---

**R**ibonucleic acid (RNA) plays important roles in many cellular processes (1, 2). To maintain their functions, secondary structures of RNA homologs are conserved across evolution (3–5). These conserved structures provide critical targets for diagnostics and treatments. Thus, there is a need for developing fast and accurate computational methods to identify structurally conserved regions.

Commonly, conserved structures involve compensatory base pair changes, where two positions in primary sequences mutate across evolution and still conserve a base pair, for instance, an AU or a CG pair replaces a GC pair in homologous sequences. These compensatory changes provide strong evidence for evolutionarily conserved structures (6–10). Meanwhile, they make it harder to align sequences when structures are unknown. Initially, the process of determining a conserved structure, termed comparative sequence analysis, was manual and required substantial insight to identify the conserved structure. A notable early achievement was the determination of the conserved tRNA secondary structure (11). Comparative analysis was also demonstrated to be 97% accurate as compared to crystal structures for ribosomal RNAs, where the models were refined carefully over time (12).

To automate comparative analysis, three distinct algorithmic approaches were developed (13, 14). The first, “joint fold-and-align” method, seeks to simultaneously predict structures and a structural alignment for two or more sequences. This was first proposed by Sankoff (15) using a dynamic programming algorithm. The major limitation of this approach is that the algorithm runs in *O*(*n*^3*k*^) against *k* sequences with the average sequence length *n*. Several software packages provide implementations of the Sankoff algorithm (16–21) that use simplifications to reduce runtime. The second, “align-then-fold” approach, is to input a sequence alignment and predict the conserved structure that can be identified across sequences in the alignment. This was described by Waterman (22), and was subsequently refined and popularized by RNAalifold (23). The third, “fold-then-align” approach, is to predict plausible structures for the sequences, and then align the structures to determine the sequence alignment and the optimal conserved structures. This was described by Waterman (24) and implemented in RNAforester (25) and MARNA (26) (*SI Appendix*, Fig. S1).

As an alternative, TurboFold II (27), an extension of TurboFold (28), provides a more computationally efficient method to align and fold sequences. Taking multiple unaligned sequences as input, TurboFold II iteratively refines alignments and structure predictions so that they conform more closely to each other and converge on conserved structures. TurboFold II is significantly more accurate than other methods (16, 18, 23, 29, 30) when tested on RNA families with known structures and alignments.

However, the cubic runtime and quadratic memory usage of TurboFold II prevent it from scaling to longer sequences such as full-length SARS-CoV-2 genomes, which contain 30,000 nucleotides; in fact, no joint-align-and-fold methods can scale to these genomes, which are the longest among RNA viruses. As a (not very principled) workaround, most existing efforts for modeling SARS-CoV-2 structures (31–36) resort to local folding methods (38, 39) with sliding windows plus a limited pairing distance, abandoning all long-range interactions, and only consider one SARS-CoV-2 genome (Fig. 1B– C), ignoring signals available in multiple homologous sequences. To address this challenge, we designed a linearized version of TurboFold II, *LinearTurboFold* (Fig. 1A), which is a global homologous folding algorithm that scales linearly with sequence length. This linear runtime makes it the first joint-fold-and-align algorithm to scale to full-length coronavirus genomes without any constraints on window size or pairing distance, taking about 13 hours to analyze a group of 25 SARS-CoV homologs. It also leads to significant improvement on secondary structure prediction accuracy as well as an alignment accuracy comparable to or higher than all benchmarks.

**Fig. 1.**
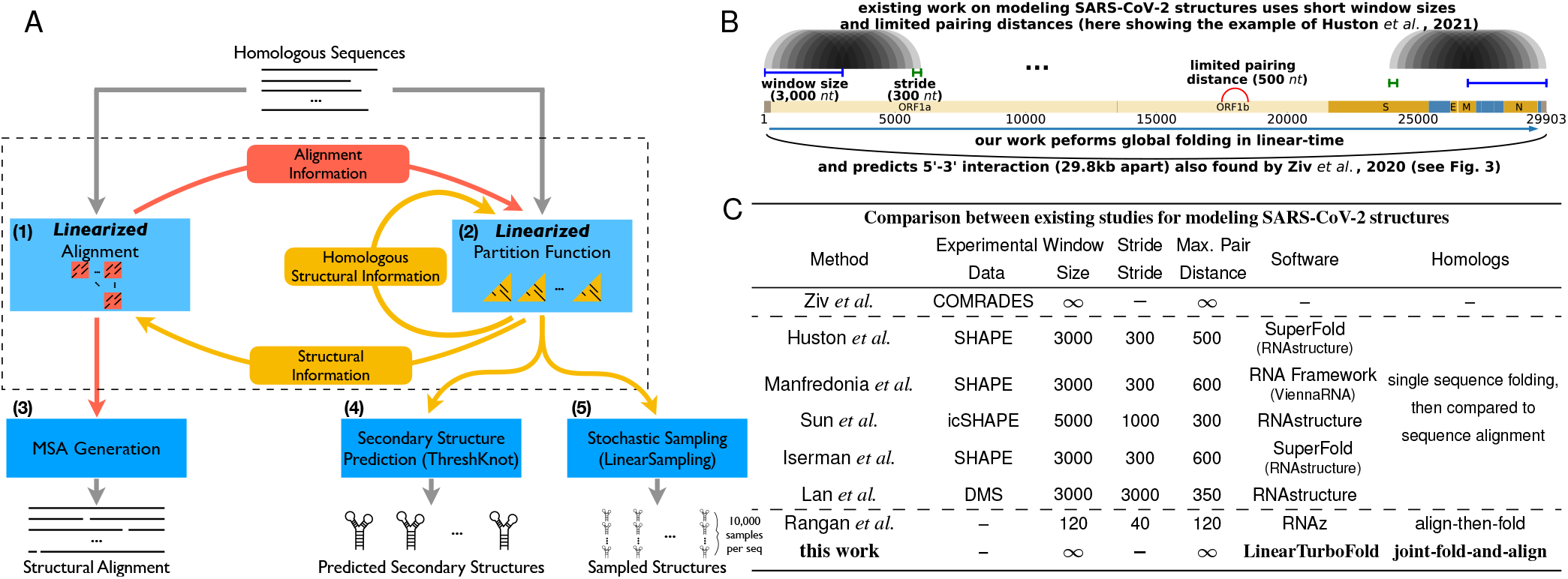
**A**: The LinearTurboFold framework. Like TurboFold II, LinearTurboFold takes multiple unaligned homologous sequences as input and outputs a secondary structures for each sequence, and a multiple sequence alignment (MSA). But unlike TurboFold II, LinearTurboFold employs two linearizations to ensure linear runtime: a *linearized* alignment computation (module **1**) to predict posterior co-incidence probabilities (red squares) for all pairs of sequences (see **Methods §1–4**), and a *linearized* partition function computation (module **2**) to estimate base-pairing probabilities (yellow triangles) for all the sequences (see **Methods §5–6**). These two modules take advantage of information from each other and iteratively refine predictions (*SI Appendix*, Fig. S2). After several iterations, module **3** generates the final multiple sequence alignments (see **Methods §7**), and module **4** predicts secondary structures. Module **5** can stochastically sample structures. **B–C**: Prior studies (31–36) (except for the purely experimental work by Ziv *et al.* (37)) used local folding methods with limited window size and maximum pairing distance. **B** shows the local folding of the SARS-CoV-2 genome by Huston *et al.*, which used a window of 3,000 *nt* that was advanced 300 *nt*. It also limited the distance between nucleotides that can base pair at 500. Some work also used homologous sequences to identify conserved structures, but they only predicted structures for one genome and utilized sequence alignments to identify mutations. By contrast, LinearTurboFold is a global folding method without any limitations on sequence length or paring distance, and it jointly folds and aligns homologs to obtain conserved structures. Consequently, LinearTurboFold can capture long-range interactions even across the whole genome (the long arc in **B** and Fig. 3).

Over a group of 25 SARS-CoV-2 and SARS-related homologous genomes, LinearTurboFold predictions are close to the canonical structures (40) and structures modeled with the aid of experimental data (32–34) for several well-studied regions. Thanks to global rather than local folding, LinearTurboFold discovers a long-range interaction involving 5’ and 3’ UTRs (~29,800 *nt* apart), which is consistent with recent purely experimental work (35), and yet is out of reach for local folding methods used by existing studies (Fig. 1B–C). In short, our *in silico* method of folding multiple homologs can achieve results similar to, and sometimes more accurate than, experimentally-guided models for one genome. Moreover, LinearTurboFold identifies conserved structures supported by compensatory mutations, which are potential targets for small molecule drugs (41) and antisense oligonucleotides (ASOs) (36). We further identify regions that are (a) sequence-level conserved, (b) at least 15 *nt* long, and (c) accessible (i.e., likely to be completely unpaired) as potential targets for ASOs (42), small interfering RNA (siRNA) (43), CRISPR-Cas13 guide RNA (gRNA) (44) and reverse transcription polymerase chain reaction (RT-PCR) primers (45). LinearTurboFold is a general technique that can also be applied to other RNA viruses (e.g., influenza, Ebola, HIV, Zika, etc.) and full-length genome studies.

## Results

The framework of LinearTurboFold has two major aspects (Fig. 1A): linearized structure-aware pairwise alignment estimation (module **1**); and linearized homolog-aware structure prediction (module **2**). LinearTurboFold iteratively refines alignments and structure predictions, specifically, updating pairwise alignment probabilities by incorporating predicted base-pairing probabilities (from module **2**) to form structural alignments, and modifying base-pairing probabilities for each sequence by integrating the structural information from homologous sequences via the estimated alignment probabilities (from module **1**) to detect conserved structures. After several iterations, LinearTurboFold generates the final multiple sequence alignment (MSA) based on the latest pairwise alignment probabilities (module **3**) and predicts secondary structures using the latest pairing probabilities (module **4**). LinearTurboFold achieves linear time regarding sequence length with two major linearized modules: our recent work LinearPartition (46) (Fig. 1A module **2**), which approximates the RNA partition function (47) and base pairing probabilities in linear time, and a novel algorithm LinearAlignment (module **1**). LinearAlignment aligns two sequences by Hidden Markov Model (HMM) in linear time by applying the same beam search heuristic (48) used by LinearPartition. Finally, LinearTurboFold assembles the secondary structure from the final base pairing probabilities using an accurate and linear-time method named ThreshKnot (49) (module **4**).

LinearTurboFold also integrates a linear-time stochastic sampling algorithm named LinearSampling (50) (module **5**), which independently samples structures according to the homolog-aware partition functions and then calculates the probability of being unpaired for regions, which is an important property in, for example, siRNA sequence design (43). Therefore, the overall end-to-end runtime of LinearTurboFold scales linearly with sequence length (**Methods §1– 7**). The number of iterations and other hyperparameters were tuned on the training set. As observed previously (27, 28), improvements after three iterations are negligible, therefore the best number of iterations is set to be three. On the testing set, it is observed that LinearTurboFold achieves the most substantial improvements in both structure prediction and MSA accuracy in the first iteration and continues to benefit from the next two iterations (*SI Appendix*, Fig. S5). which is consistent with the observation on the training set. After approximately three iterations, both structure prediction and MSA accuracies remain stable with small fluctuations. To better demonstrate the improvement in each iteration, we visualized both base-pairing probabilities and alignment co-incidence probabilities from LinearTurboFold for a group of five tRNAs across iterations (*SI Appendix*, Fig. S6–S7).

### Scalability and Accuracy

To evaluate the efficiency of LinearTurboFold against the sequence length, we collected a dataset consisting of seven families of RNAs with sequence length ranging from 210 *nt* to 30,000 *nt*, including five families from the RNAStrAlign dataset (27) plus 23S ribosomal RNA, HIV genomes and SARS-CoV genomes, and the calculation for each family uses five homologous sequences (**Methods §8**). Fig. 2A compares the running times of LinearTurboFold with TurboFold II and two Sankoff-style simultaneous folding and alignment algorithms, LocARNA and MXSCARNA. Clearly, LinearTurboFold scales linearly with sequence length *n*, and is substantially faster than other algorithms, which scale superlinearly. The linearization in LinearTurboFold brought orders of magnitude speedup over the cubic-time TurboFold II, taking only 12 minutes on the HIV family (average length 9,686 *nt*) while TurboFold II takes 3.1 days (372× speedup). More importantly, LinearTurboFold takes only 40 minutes on five SARS-CoV sequences while all other benchmarks fail to scale. Regarding the memory usage (Fig. 2B), LinearTurboFold costs linear memory space with sequence length, while other benchmarks use quadratic or more memory. In Fig. 2C–D, we also demonstrate that the runtime and memory usage against the number of homologs using sets of 16S rRNAs about 1,500 *nt* in length. The apparent complexity of LinearTurboFold against the group size *k* is higher than that of TurboFold II because the runtime of the latter is *O*(*kn*^3^ + *k*^2^*n*^2^) and is dominated by the *O*(*kn*^3^) partition function calculation, thus scaling *O*(*k*^1.4^) empirically. By contrast, LinearTurboFold linearizes both partition function and alignment modules, so its overall runtime becomes *O*(*kn* + *k*^2^*n*) and is instead dominated by the *O*(*k*^2^*n*) alignment module, therefore scaling *O*(*k*^2^) in practice. A similar analysis holds for memory usage (Fig. 2E).^*^

**Fig. 2.**
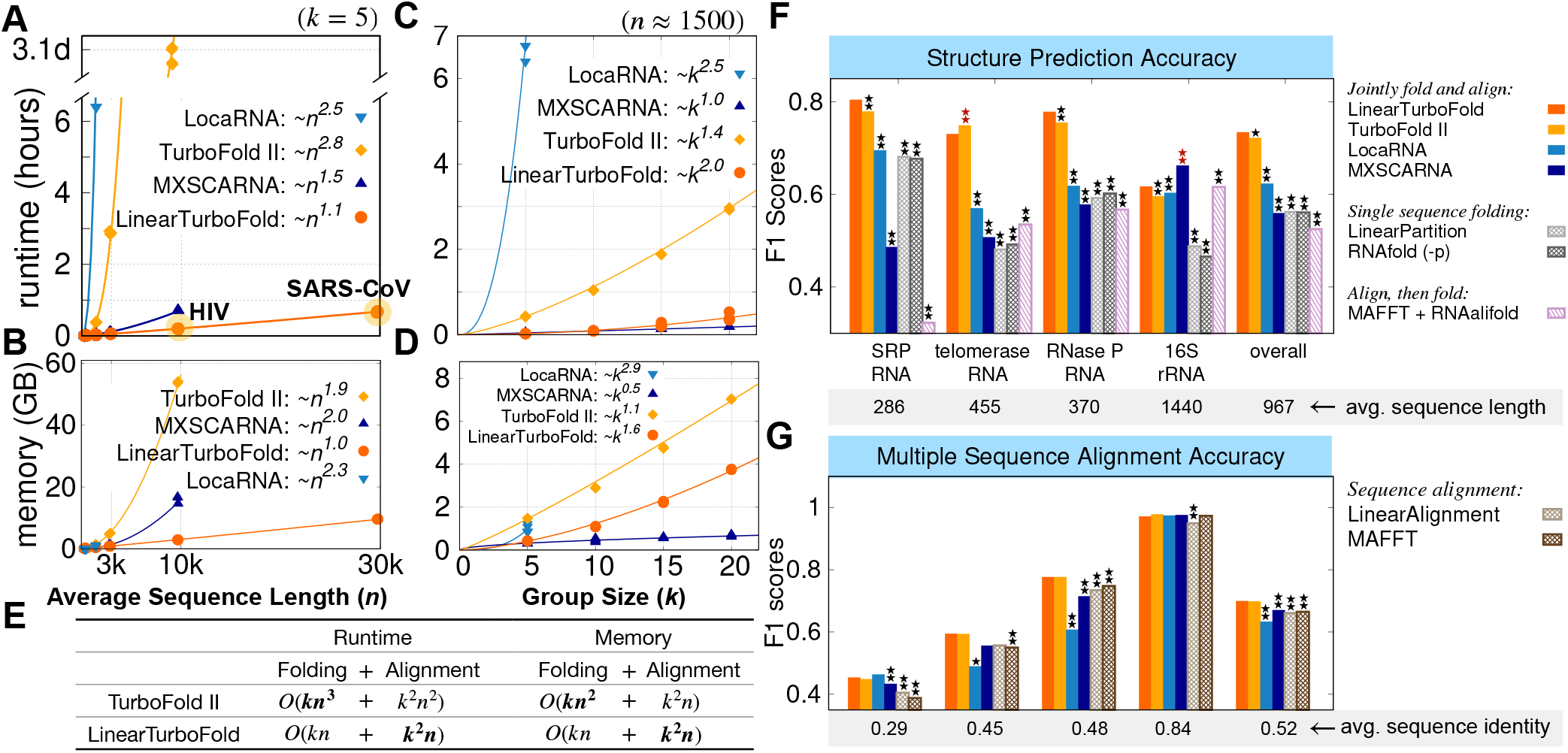
End-to-end Scalability and Accuracy Comparisons. **A–B**: End-to-end runtime and memory usage comparisons between benchmarks and LinearTurboFold against the sequence length. LinearTurboFold uses beam size 100 in both partition function and HMM alignment calculation with three iterations to run all groups of data. **C–D**: End-to-end runtime and memory usage comparisons against the group size. LinearTurboFold is the first joint-fold-and-align algorithm to scale to full-length coronavirus genomes ( 30,000 *nt*) due to its linear runtime. **E**: The runtime and space complexity comparisons between TurboFold II and LinearTurboFold. The dominating terms are in bold. **F–G**: The F1 accuracy scores of the structure prediction and multiple sequence alignment (*SI Appendix*, Tab. S1). LocARNA and MXSCARNA are Sankoff-style simultaneous folding and alignment algorithms for homologous sequences. As negative controls, LinearPartition and Vienna RNAfold-predicted structures for each sequence separately; LinearAlignment and MAFFT generated sequence-level alignments; RNAalifold folded pre-aligned sequences (e.g., from MAFFT) and predicted conserved structures. Statistical significances (two-tailed permutation test) between the benchmarks and LinearTurboFold are marked with one star (*\**) on the top of the corresponding bars if *p* < 0.05 or two stars 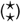 if *p* < 0.01. The benchmarks whose accuracies are significantly lower than LinearTurboFold are annotated with black stars, while benchmarks higher than LinearTurboFold are marked with dark red stars. Overall, on structure prediction, LinearTurboFold achieves significantly higher accuracy than all evaluated benchmarks, and on multiple sequence alignment, it achieves accuracies comparable to TurboFold II and significantly higher than other methods (*SI Appendix*, Tab. S1).

We next compare the accuracies of secondary structure prediction and MSA between LinearTurboFold and several benchmark methods (**Methods §9**). Besides Sankoff-style LocARNA and MXSCARNA, we also consider three types of negative controls: (a) single sequence folding (partition function-based): Vienna RNAfold (39) (-p mode) and LinearPartition; (b) sequence-only alignment: MAFFT (29) and LinearAlignment (a standalone version of the alignment method developed for this work but without structural information); and (c) an align-then-fold method that predicts consensus structures from MSAs (*SI Appendix*, Fig. S1): MAFFT + RNAalifold (23).

For secondary structure prediction, LinearTurboFold, TurboFold II and LocARNA achieve higher F1 scores than single sequence folding methods (Vienna RNAfold and LinearPartition) (Fig. 2F), which demonstrates folding with homology information performs better than folding sequences separately. Overall, LinearTurboFold performs significantly better than all the other benchmarks on structure prediction. For the accuracy of MSAs (Fig. 2G), the structural alignments from LinearTurboFold obtain higher accuracies than sequence-only alignments (LinearAlignment and MAFFT) on all four families, especially for families with low sequence identity. On average, LinearTurboFold performs comparably with TurboFold II and significantly better than other benchmarks on alignments. We also note that the structure prediction accuracy of the align-then-fold approach (MAFFT + RNAalifold) depends heavily on the alignment accuracy, and is the worst when the sequence identity is low (e.g., SRP RNA) and the best when the sequence identity is high (e.g., 16S rRNA) (Fig. 2F–G).

### Highly Conserved Structures in SARS-CoV-2 and SARS-related Betacoronaviruses

RNA sequences with conserved secondary structures play vital biological roles and provide potential targets. The current COVID-19 outbreak raises an emergent requirement of identifying potential targets for diagnostics and therapeutics. Given the strong scalability and high accuracy, we used LinearTurboFold on a group of full-length SARS-CoV-2 and SARS-related (SARSr) genomes to obtain global structures and identify highly conserved structural regions.

We used a greedy algorithm to select the 16 most diverse genomes from all the valid SARS-CoV-2 genomes submitted to the Global Initiative on Sharing Avian Influenza Data (GISAID) (52) up to December 2020 (**Methods §11**). We further extended the group by adding 9 SARS-related homologous genomes (5 human SARS-CoV-1 and 4 bat coronaviruses) (53). In total, we built a dataset of 25 fulllength genomes consisting of 16 SARS-CoV-2 and 9 SARS-related sequences (*SI Appendix*, Fig. S9).The average pairwise sequence identities of the 16 SARS-CoV-2 and the total 25 genomes are 99.9% and 89.6%, respectively. LinearTurboFold takes about 13 hours and 43 GB on the 25 genomes.

To evaluate the reliability of LinearTurboFold predictions, we first compare them with the Huston *et al.*’s SHAPE-guided models (32) for regions with well-characterized structures across betacoronaviruses. For the extended 5’ and 3’ untranslated regions (UTRs), LinearTurboFold’s predictions are close to the SHAPE-guided structures (Fig. 3A–B), i.e., both identify the stem-loops (SLs) 1–2 and 4–7 in the extended 5’ UTR, and the bulged stem-loop (BSL), SL1, and a long bulge stem for the hypervariable region (HVR) including the stem-loop II-like motif (S2M) in the 3’ UTR. Interestingly, in our model, the high unpaired probability of the stem in the SL4b indicates the possibility of being single-stranded as an alternative structure, which is supported by experimental studies (33, 36). In addition, the compensatory mutations LinearTurboFold found in UTRs strongly support the evolutionary conservation of structures (Fig. 3A).

**Fig. 3.**
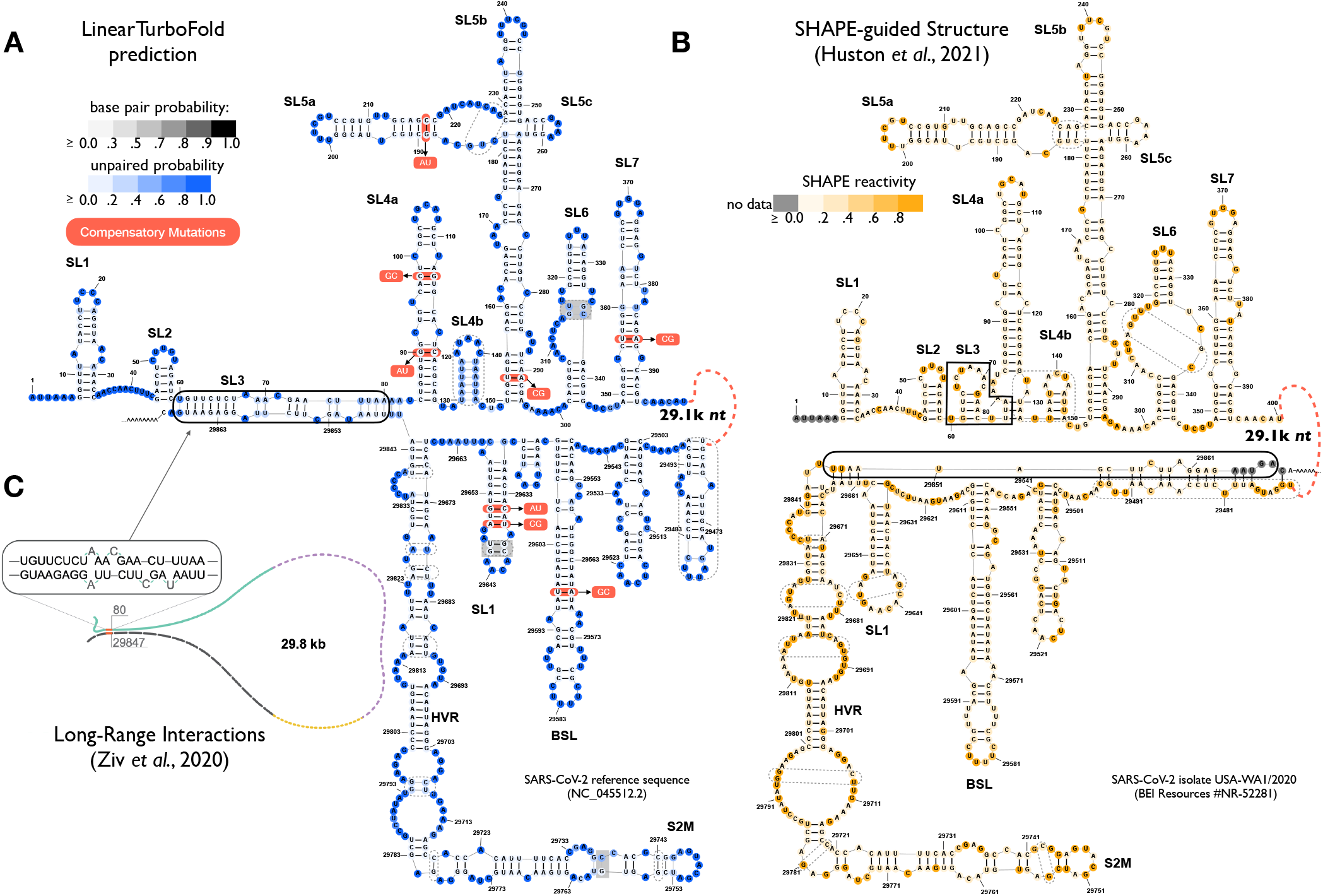
Secondary structures predictions of SARS-CoV-2 extended 5’ and 3’ UTRs. **A**: LinearTurboFold prediction.The nucleotides and base pairs are colored by unpaired probabilities and base-pairing probabilities, respectively. The compensatory mutations extracted by LinearTurboFold are annotated with alternative pairs in red boxes (see *SI Appendix*, Tab. S2 for more fully conserved pairs with co-variational changes). **B**: SHAPE-guided model by Huston *et al.* (32) (window size 3000 *nt* sliding by 300 *nt* with maximum pairing distance 500 *nt*). The nucleotides are colored by SHAPE reactivities. Dashed boxes enclose the different structures between **A** and **B**. Our model is close to Huston *et al.*’s, but the major difference is that LinearTurboFold predicts the end-to-end pairs involving 5’ and 3’ UTRs (solid box in **A**), which is *exactly* the same interaction detected by Ziv *et al.* using the COMRADES experimental technique (37) (**C**). Such long-range interactions cannot be captured by the local folding methods used by prior experimentally-guided models (Fig. 1B). The similarity between models A and B as well as the exact agreement between A and C show that our *in silico* method of folding multiple homologs can achieve results similar to, if not more accurate than, experimentally-guided single-genome prediction. As negative controls (*SI Appendix*, Fig. S10), the align-then-fold (RNAalifold) method cannot predict such long-range interactions. Although the single sequence folding algorithm (LinearPartition) predicts a long-range 5’-3’ interaction, the positions are not the same as the LinearTurboFold model and Ziv *et al.*’s experimental result.

The most important difference between LinearTurboFold’s prediction and Huston *et al.*’s experimentally-guided model is that LinearTurboFold discovers an end-to-end interaction (29.8 kilobases apart) between the 5’ UTR (SL3, 60-82 *nt*) and the 3’ UTR (final region, 29845-29868 *nt*), which fold locally by themselves in Huston *et al.*’s model. Interestingly, this 5’-3’ interaction matches *exactly* with the one discovered by the purely experimental work of Ziv *et al.* (37) using the COMRADES technique to capture long-range base-pairing interactions (Fig. 3C). These end-to-end interactions have been well established by theoretical and experimental studies (54–56) to be common in natural RNAs, but are far beyond the reaches of local folding methods used in existing studies on SARS-CoV-2 secondary structures (32–35). By contrast, LinearTurboFold predicts secondary structures globally without any limit on window size or base-pairing distance, enabling it to discover long-distance interactions across the whole genome. The similarity between our predictions and the experimental work shows that our *in silico* method of folding multiple homologs can achieve results similar to, if not more accurate than, those experimentally-guided single-genome prediction.

LinearTurboFold can model these end-to-end interactions thanks to three ingredients: (a) linearization, (b) LinearPartition’s better modeling power on long sequences and long-range pairs, and (c) homologous folding and soft alignment. Linearization not only enables LinearTurboFold to scale to longer sequences, but also improves the accuracy of modeling long-range interactions benefiting from LinearPartition (46). In addition, homologous folding is also crucial. We observed that LinearPartition can model the same end-to-end interactions detected by Ziv *et al.* for 8 out of 25 sequences (4 out of 16 SARS-CoV-2 and 4 out of 9 SARS-related sequences; see *SI Appendix*, Fig. S12A and the left column of Fig. S13). For the other sequences, however, LinearPartition either cannot predict end-to-end interactions or predicts them in the wrong locations. On the other hand, LinearTurboFold propagates the correct structural information from those eight sequences to other homologs, resulting in all SARS-CoV-2 sequences having the same end-to-end pairs (*SI Appendix*, Fig. S12B and the right column of Fig. S13). By contrast, the align-then-fold approach (MAFFT + RNAalifold), which relies on the input hard alignment and predicts one single consensus structure for all homologs, fails to predict such long-range interactions (*SI Appendix*, Fig. S10B).

The frameshifiting stimulation element (FSE) is another well-characterized region. For an extended FSE region, the LinearTurboFold prediction consists of two substructures (Fig. 4A): the 5’ part includes an attenuator hairpin and a stem, which are connected by a long internal loop (16 *nt*) including the slippery site, and the 3’ part includes three stem loops. We observe that our predicted structure of the 5’ part is consistent with experimentally-guided models (32, 33, 35) (Fig. 4B–D). In the attenuator hairpin, the small internal loop motif (UU) was previously selected as a small molecule binder that stabilizes the folded state of the attenuator hairpin and impairs frameshifting (41). For the long internal loop including the slippery site, we will show in the next section that it is both highly accessible and conserved (Fig. 5), which makes it a perfect candidate for drug design. For the 3’ region of the FSE, LinearTurboFold successfully predicts stems 1–2 (but misses stem 3) of the canonical three-stem pseudoknot (40) (Fig. 4E). Our prediction is closer to the canonical structure compared to the experimentally-guided models (32, 33, 35) (Fig. 4B–D); one such model (Fig. 4B) identified the pseudoknot (stem 3) but with an open stem 2. Note that all these experimentally-guided models for the FSE region were estimated for specific local regions. As a result, the models are sensitive to the context and region boundaries (32, 35, 57) (see *SI Appendix*, S11D–F for alternative structures of Fig. 4B–D with different regions). LinearTurboFold, by contrast, does not suffer from this problem by virtue of global folding without local windows. Besides SARS-CoV-2, we notice that the estimated structure of the SARS-CoV-1 reference sequence (Fig. 4F) from LinearTurboFold is similar to SARS-CoV-2 (Fig. 4A), which is consistent with the observation that the structure of the FSE region is highly conserved among betacoronaviruses (40). Finally, as negative controls, both the single sequence folding algorithm (LinearPartition in Fig. 4G) and the align-then-fold method (RNAalifold in *SI Appendix*, Fig. S11G) predict quite different structures compared with the LinearTurboFold prediction (Fig. 4A) (39%/61% of pairs from the LinearTurboFold model are not found by LinearPartition/RNAalifold).

**Fig. 4.**
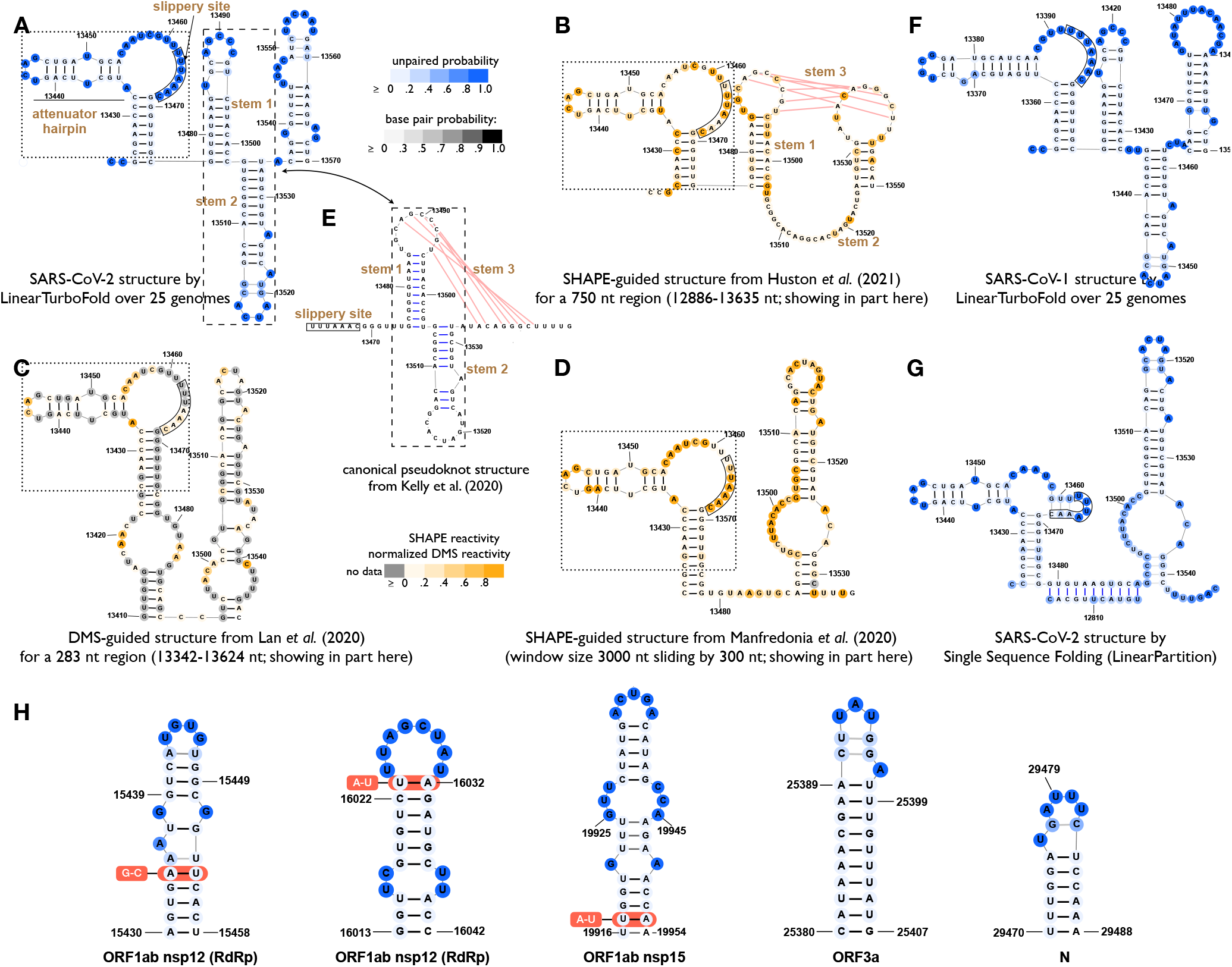
**A–D**: Secondary structure predictions of SARS-CoV-2 extended frameshifting stimulation element (FSE) region (13425–13545 *nt*). **A**: LinearTurboFold prediction. **B–D**: Experimentally-guided predictions from the literature (32, 33, 35), which are sensitive to the context and region boundaries due to the use of local folding methods (*SI Appendix*, Fig. S11). **E**: The canonical pseudoknot structure by the comparative analysis between SARS-CoV-1 and SARS-CoV-2 genomes (40). For the 5’ region of the FSE shown in dotted boxes (attenuator hairpin, internal loop with slippery site, and a stem), the LinearTurboFold prediction (A) is consistent with B–D; for the 3’ region of the FSE shown in dashed boxes, our prediction (predicting stems 1–2 but missing 3) is closer to the canonical structure in E compared to B–D. **F**: LinearTurboFold prediction on SARS-CoV-1. **G**: Single sequence folding algorithm (LinearPartition) prediction on SARS-CoV-2, which is quite different from LinearTurboFold’s. As another negative control, the align-then-fold method (RNAalifold) predicts a rather dissimilar structure (*SI Appendix*, Fig. S11G). **H**: Five examples from 59 fully conserved structures among 25 genomes (*SI Appendix*, Tab. S3), 26 of which are novel compared with prior work (31, 32).

**Fig. 5.**
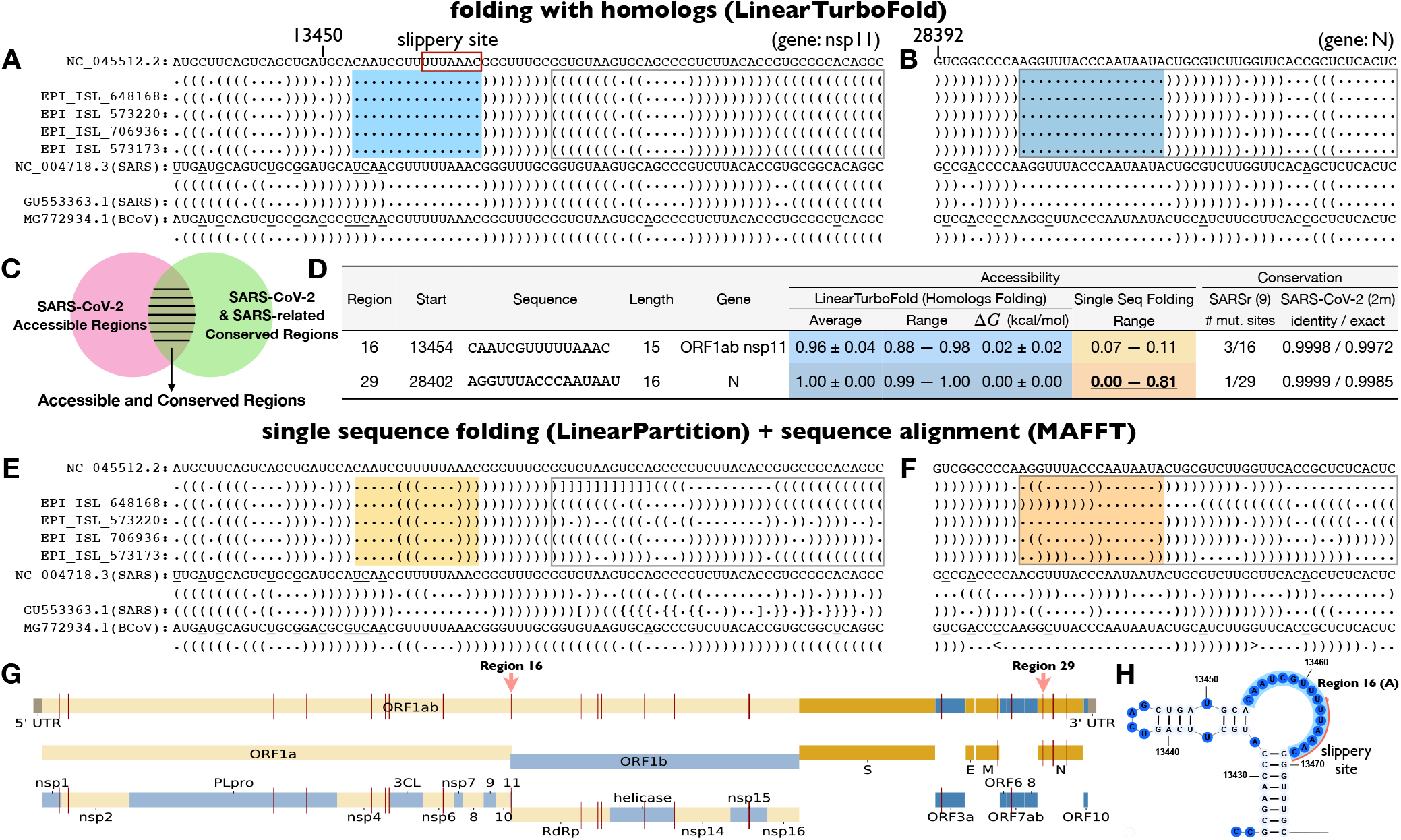
An illustration of accessible and conserved regions that LinearTurboFold identifies. **A–B**: Identified structurally-conserved accessible regions by LinearTurboFold with the help of considering alignment and folding simultaneously. The regions at least 15 *nt* long with accessibility of at least 0.5 among all the 16 SARS-CoV-2 genomes are shaded on blue background. Structures are encoded in dot-bracket notation. “(” and “)” indicates nucleotides pairing in the 3’ and 5’ direction, respectively. “.” indicates an unpaired nucleotide. The positions with mutations compared to the SARS-CoV-2 reference sequence among three different subfamilies (SARS-CoV-2, SARS-CoV-1 and BCoV) are underlined. **C**: Accessible and conserved regions are not only *accessible* among SARS-CoV-2 genomes (pink circle) but also *conserved* (at sequence level) among both SARS-CoV-2 and SARS-related genomes (green circle). **D**: Two examples out of 33 accessible and conserved regions found by LinearTurboFold. Region 16 and Region 29 correspond to the accessible regions in **A** and **B**, respectively. Region 16 is also the long internal loop including the slippery site in the FSE region (**H**). The conservation of these regions on 9 SARS-related genomes is the number of mutated sites. The conservation on the ∼2M SARS-CoV-2 dataset is shown in both average sequence identity with the reference sequence and the percentage of exact matches, respectively. **E–F**: Single sequence folding algorithms predict greatly different structures even if the sequence identities are high (grey boxes). These two regions, fully conserved among SARS-CoV-2 genomes, still fold into different structures due to mutations outside the regions. **G**: The positions of these 33 regions (red bars) across the whole genome (*SI Appendix*, Tab. S5). All the accessible and conserved regions are potential targets for siRNAs, ASOs, CRISPR-Cas13 gRNAs and RT-PCR primers.

In addition to the well-studied UTRs and FSE regions, LinearTurboFold discovers 50 conserved structures with identical structures among 25 genomes, and 26 regions are novel compared to previous studies (31, 32) (Fig. 4H, *SI Appendix*, Tab. S3). These novel structures are potential targets for small-molecule drugs (41) and ASOs (36, 58). LinearTurboFold also recovers fully conserved base pairs with compensatory mutations (*SI Appendix*, Tab. S2), which imply highly conserved structural regions whose functions might not have been explored. We provide the complete multiple sequence alignment and predicted structures for 25 genomes from LinearTurboFold (Dataset S1; see *SI Appendix*, Fig. S14 for the format).

### Highly Accessible and Conserved Regions in SARS-CoV-2 and SARS-related Betacoronaviruses

Studies show that the siRNA silencing efficiency, ASO inhibitory efficacy, CRISPR-Cas13 knockdown efficiency, and RT-PCR primer binding efficiency, all correlate with the target region’s *accessibility* (43–45, 59), which is the probability of a target site being fully unpaired. However, most existing work for designing siRNAs, ASOs, CRISPR-Cas13 gRNAs, and RT-PCR primers does not take this feature into consideration (60, 61) (*SI Appendix*, Tab. S4). Here LinearTurboFold is able to provide more principled design candidates by identifying accessible regions of the target genome. In addition to accessibility, the emerging variants around the world reduce effectiveness of existing vaccines and test kits (*SI Appendix*, Tab. S4), which indicates sequence conservation is another critical aspect for therapeutic and diagnostic design. LinearTurboFold, being a tool for both structural alignment and homologous folding, can identify regions that are both (sequence-wise) conserved and (structurally) accessible, and it takes advantage of not only SARS-CoV-2 variants but also homologous sequences, e.g., SARS-CoV-1 and bat coronavirus genomes, to identify conserved regions from historical and evolutionary perspectives.

To get unstructured regions, Rangan *et al.* (31) imposed a threshold on unpaired probability of each position, which is a crude approximation because the probabilities are not independent of each other. By contrast, the widely-used stochastic sampling algorithm (50, 62) builds a representative ensemble of structures by sampling independent secondary structures according to their probabilities in the Boltzmann distribution. Thus the accessibility for a region can be approximated as the fraction of sampled structures in which the region is single-stranded. LinearTurboFold utilized LinearSampling (50) to generate 10,000 independent structures for each genome according to the modified partition functions after the iterative refinement (Fig. 1A module **5**), and calculated accessibilities for regions at least 15 *nt* long. We then define *accessible regions* that are with at least 0.5 accessibility among all 16 SARS-CoV-2 genomes (Fig. 5A–B). We also measure the free energy to open a target region [*i, j*] (63), notated: Δ*G*_u_[*i, j*] = − *RT*(log *Z*_u_[*i, j*] − log *Z*) = − *RT* log *P*_u_[*i, j*] where *Z* is the partition function which sums up the equilibrium constants of all possible secondary structures, *Z*_u_[*i, j*] is the partition function over all structures in which the region [*i, j*] is fully unpaired, *R* is the universal gas constant and *T* is the thermodynamic temperature. Therefore *P*_u_[*i, j*] is the unpaired probability of the target region and can be approximated via sampling by *s*0/*s*, where *s* is the sample size and *s*0 is the number of samples in which the target region is single-stranded. The regions whose free energy changes are close to zero need less free energy to open, thus more accessible to bind with siRNAs, ASOs, CRISPR-Cas13 gRNAs and RT-PCR primers.

Next, to identify *conserved regions* that are highly conserved among both SARS-CoV-2 and SARS-related genomes, we require that these regions contain at most three mutated sites on the 9 SARS-related genomes compared to the SARS-CoV-2 reference sequence because historically conserved sites are also unlikely to change in the future (64), and the average sequence identity with reference sequence over a large SARS-CoV-2 dataset is at least 0.999 (here we use a dataset of 2M SARS-CoV-2 genomes submitted to GISAID up to June 30, 2021^†^; see **Methods §11**). Finally, we identified 33 *accessible and conserved regions* (Fig. 5G and *SI Appendix*, Tab. S5), which are not only structurally accessible among SARS-CoV-2 genomes but also highly conserved among SARS-CoV-2 and SARS-related genomes (Fig. 5C). Because the specificity is also a key factor influencing siRNA efficiency (65), we used BLAST against the human transcript dataset for these regions (*SI Appendix*, Tab. S5). Finally, we also listed the GC content of each region. Among these regions, region 16 corresponds to the internal loop containing the slippery site in the extended FSE region, and it is conserved at both structural and sequence levels (Fig. 5D and 5H). Besides SARS-CoV-2 genomes, the SARS-related genomes such as the SARS-CoV-1 reference sequence (NC_004718.3) and a bat coronavirus (BCoV, MG772934.1) also form similar structures around the slippery site (Fig. 5A). By removing the constraint of conservation on SARS-related genomes, we identified 38 additional candidate regions (*SI Appendix*, Tab. S6) that are accessible but only highly conserved on SARS-CoV-2 variants.

We also designed a negative control by analyzing the SARS-CoV-2 reference sequence alone using LinearSampling, which can also predict accessible regions. However, these regions are not structurally conserved among the other 15 SARS-CoV-2 genomes, resulting in vastly different accessibilities, except for one region in the M gene (*SI Appendix*, Tab. S7). The reason for this difference is that, even with a high sequence identity (over 99.9%), single sequence folding algorithms still predict greatly dissimilar structures for the SARS-CoV-2 genomes (Fig. 5E–F). Both regions (in nsp11 and N genes) are fully conserved among the 16 SARS-CoV-2 genomes, yet they still fold into vastly different structures due to mutations outside the regions; as a result, the accessibilities are either low (nsp11) or in a wide range (N) (Fig. 5D). Conversely, addressing this by folding each sequence with proclivity of base pairing inferred from all homologous sequences, LinearTurboFold structure predictions are more consistent with each other and thus can detect conserved structures (Fig. 5A–B).

## Discussion

The constant emergence of new SARS-CoV-2 variants is reducing the effectiveness of exiting vaccines and test kits. To cope with this issue, there is an urgent need to identify conserved structures as promising targets for therapeutics and diagnostics that would work in spite of current and future mutations. Here we presented LinearTurboFold, an end-to-end linear-time algorithm for structural alignment and conserved structure prediction of RNA homologs, which is the first joint-fold-and-align algorithm to scale to full-length SARS-CoV-2 genomes without imposing any constraints on base-pairing distance. We also demonstrate that LinearTurboFold leads to significant improvement on secondary structure prediction accuracy as well as an alignment accuracy comparable to or higher than all benchmarks.

Unlike existing work on SARS-CoV-2 using local folding and single-sequence folding workarounds, LinearTurboFold enables unprecedented global structural analysis on SARS-CoV-2 genomes; in particular, it can capture long-range interactions, especially the one between 5’ and 3’ UTRs across the whole genome, which matches perfectly with a recent purely experiment work. Over a group of SARS-CoV-2 and SARS-related homologs, LinearTurboFold identifies not only conserved structures supported by compensatory mutations and experimental studies, but also accessible and conserved regions as vital targets for designing efficient small-molecule drugs, siRNAs, ASOs, CRISPR-Cas13 gRNAs and RT-PCR primers. LinearTurboFold is widely applicable to the analysis of other RNA viruses (influenza, Ebola, HIV, Zika, etc.) and full-length genome analysis.

## Methods

### §1 Pairwise Hidden Markov Model

We use a pairwise Hidden Markov Model (pair-HMM) to align two sequences (51, 66). The model includes three actions (*h*): aligning two nucleotides from two sequences (ALN), inserting a nucleotide in the first sequence without a corresponding nucleotide in the other sequence (INS1), and a nucleotide insertion in the second sequence without a corresponding nucleotide in the first sequence (INS2). We then define 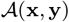 as a set of all the possible alignments for the two sequences, and one alignment 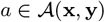 as a sequence of steps (*h, i, j*) with *m* + 2 steps, where (*h, i, j*) means an alignment step at the position pair (*i, j*) by the action *h*. Thus, for the *l*th step *a*_*l*_ = (*h*_*l*_, *i*_*l*_, *j*_*l*_) *∈ a*, the values of *i*_*l*_ and *j*_*l*_ depend on the action *h*_*l*_ and the positions*i*_*l*−1_ and *j*_*l*−1_ of *a*_*l*−1_:

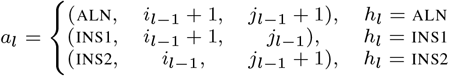

with (ALN, 0, 0) as the first step, and (ALN, |**x**| + 1, |**y**| + 1) as the last one. For two sequences {ACAAGU, AACUG}, one possible alignment {−ACAAGU, AAC− −UG} can be specified as {(ALN, 0, 0) → (INS2, 0, 1) → (ALN, 1, 2) → (ALN, 2, 3) → (INS1, 3, 3) → (INS1, 4, 3) → (ALN, 5, 4) → (ALN, 6, 5) → (ALN, 7, 6)}, where a gap symbol (−) represents a nucleotide insertion in the other sequence at the corresponding position (*SI Appendix*, Tab. S3). The action *h_l_* in each step (*h*_*l*_, *i*_*l*_, *j*_*l*_) corresponds to a line segment starting from the previous node (*i*_*l*−1_, *j*_*l*−1_) and stopping at the node (*i*_*l*_, *j*_*l*_). Thus the line segment is horizontal, vertical or diagonal towards the top-right corner when *h*_*l*_ is INS1, INS2 or ALN, respectively (*SI Appendix*, Tab. S3).

We initialize the first step with the state _ALN_ of probability 1, thus *p*_*π*_(_ALN_) = 1. *p*_t_(*h*_2_ | *h*_1_) is the transition probability from the state *h*_1_ to *h*_2_, and *p*_e_((*c*_1_, *c*_2_) | *h*_1_) is the probability of the state *h*_1_ emitting a character pair (*c*_1_, *c*_2_) with values from {A, G, C, U, −}. Both the emission and transition probabilities were taken from TurboFold II. The function *e*() yields a character pair based on *a*_*l*_ and the nucleotides of two sequences:

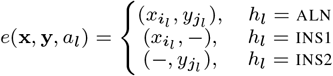

where *x*_*i*_ and *y*_*i*_ are the *i*th and *j*th nucleotides of sequences **x** and **y**, respectively. Note that the first step *a*_0_ = (ALN, 0, 0) and the last *a*_*m*+1_ = (ALN, |**x**| + 1, |**y**| + 1) do not have emissions.

We denote forward probability 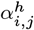 encompassing the probability of the partial alignments of **x** and **y** up to positions *i* and *j*, and all the alignments that go through the step (*h, i, j*):

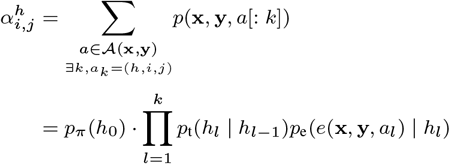

where *a*[: *k*] indicates the partial alignments from the starting node up to the *k*th step and *a*_*k*_ = (*h, i, j*). For instance, 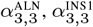 and 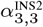 corresponds to the region circled by the blue dashed lines (*SI Appendix*, Tab. S3B, C and D). Similarly, the backward probability 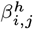 assembles the probability of partial alignments *a*[*k* + 1 :] from the (*k* + 1)th step up to the end one

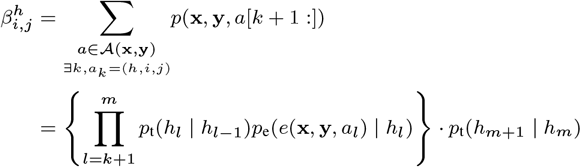

For example, 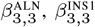 and 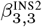 are the regions circled by the yellow dashed line (*SI Appendix*, Tab. S3B, C and D). Thus, the probability of observing two sequences *p*(**x**, **y**) is 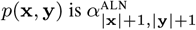 or 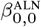.

### §2 Posterior Co-incidence Probability Computation

Nucleotide positions *i* and *j* in two sequences x and y are said to be *co-incident* (notated as *i ∼ j*) in an alignment *a* if the alignment path goes through the node (*i, j*) (51). Since the node (*i, j*) is reachable by three actions 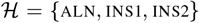, the co-incidence probability for a position pair (*i, j*) given two sequences is:

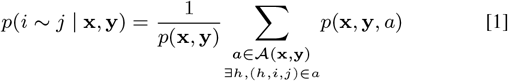

where *p*(**x**, **y**, *a*) is the probability of two sequences with the alignment *a*, and *p*(**x**, **y**) is the probability of observing two sequences, which is the sum of probability of all the possible alignments:

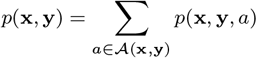

The co-incidence probability for positions *i* and *j* (Equation 1) can be computed by:

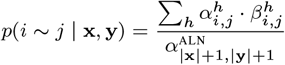

### §3 LinearAlignment

Unlike a previous method (51) that fills out all the nodes in the alignment matrix by columns (*SI Appendix*, Fig. S3). LinearAlignment scans the matrix based on the *step count s*, which is the sum value of *i* and *j* (*s* = *i* + *j*) for the partial alignments of **x**_[1,*i*]_ and **y**_[1,*j*]_. As shown in the pseudocode (*SI Appendix*, Fig. S4), the forward phase starts from the node (0, 0) in the state _ALN_ of probability 1, then iterates the step count *s* from 0 to |**x**| + |**y**| − 1. For each step count *s* with a specific state *h* from 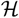, we first collect all the nodes (*i, j*) with the step count *s* with 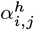 existing, which means the position pair (*i, j*) has been visited via the state *h* before. Then each node makes transitions to next nodes by there states, ans updates the tcorresponding forward probabilities 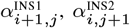 and 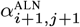, respectively.

The current alignment algorithm is still an exhaustive-search algorithm and costs quadratic time and space for all the |**x**| × |**y**| nodes. To reduce the runtime, LinearAlignment uses the beam search heuristic algorithm (48) and keeps a limited number of promising nodes at each step. For each step count *s* with a state *h*, LinearAlignment applies the beam search method first over *B*(*s, h*), which is the collection of all the nodes (*i, j*) with step count *s* and the presence of 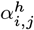 (*SI Appendix*, Fig. S4 line 6).This algorithm only saves the top *b*_*aln*_ nodes with the highest forward scores in *B*(*s, h*), and these are subsequently allowed to make transitions to the next states. Here *b*_*aln*_ is a user-specified beam size and the default value is 100. In total, *O*(*b_aln_n*) nodes survive because the length of *s* is |**x**| + |**y**| and each step count keeps *b*_*aln*_ nodes. For simplicity, we show the topological order and the beam search method with alignment examples (*SI Appendix*, Fig. S3A), while the forwardbackward algorithm adopts the same idea by summing the probabilities of all the possible alignments.

After the forward phase, the backward phase (*SI Appendix*, Fig. S4) performs in linear time to calculate the co-incidence probabilities automatically because only a linear number of nodes in *B*(*s, h*) are stored. Thus by pruning low-scoring candidates at each step in the forward algorithm, we reduce the runtime from *O*(*n*^2^) to *O*(*b_aln_n*) for aligning two sequences. For *k* input homologous sequences, LinearTurboFold computes posterior co-incidence probabilities for each pair of sequences by LinearAlignment, which costs *O*(*k*^2^*b*_*aln*_*n*) runtime in total.

### §4 Match Scores Computation and Modified LinearAlignment

To encourage the pairwise alignment conforming with estimated secondary structures, LinearTurboFold predicts structural alignments by incorporating the secondary structural conformation. PMcomp (67) first proposed the match score to measure the structural similarity for position pairs between a pair of sequences, and TurboFold II adapts it as a prior. Based on the base pair probabilities *P*_x_(*i, j*) estimated from the partition function for a sequence x, a position *i* could be paired with bases upstream, downstream or unpaired, with corresponding probability 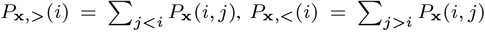 and *P*_x,*o*_(*i*) = 1 − *P*_x,>_(*i*) − *P*_x,<_(*i*), respectively. The match score *m*_x,y_(*i, j*) for two positions *i* and *j* from two sequences x and y is based on the probabilities of these three structural propensities from the last iteration (*t* − 1):

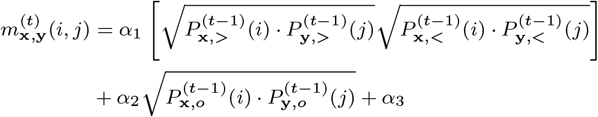

where *α*_1_, *α*_2_ and *α*_3_ are weight parameters trained in TurboFold II. The forward-backward phrases integrate the match score as a prior when aligning two nucleotides (*SI Appendix*, Fig. S4 line 10 and line 12).

TurboFold II separately pre-computes match scores for all the *O*(*n*^2^) position pairs for pairs of sequences before the HMM alignment calculation. However, only a linear number of pairs *O*(*b*_*aln*_*n*) survive after applying the beam pruning in LinearAlignment. To reduce redundant time and space usage, LinearTurboFold calculates the corresponding match scores for co-incident pairs when they are first visited in LinearAlignment. Overall, for *k* homologous sequences, LinearTurboFold reduces the runtime of the whole module of pairwise posterior co-incidence probability computation from *O*(*k*^2^*n*^2^) to *O*(*k*^2^*b*_*aln*_*n*) by applying the beam search heuristic to the pairwise HMM alignment, and only calculating the match scores for position pairs that are needed.

### §5 Extrinsic Information Calculation

To update partition functions for each sequence with the structural information from homologs, TurboFold (28) introduces *extrinsic information* to model the the proclivity for base pairing induced from the other sequences in the input set 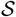. The extrinsic information *e*x(*i, j*) for a base pair (*i, j*) in the sequence x maps the estimated base pairing probabilities of other sequences to the target sequence via the co-incident nucleotides between each pair of sequences:

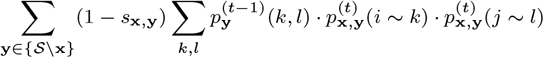

where 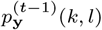 is the base pair probability for a base pair (*k, l*) in the sequence **y** from (*t* − 1)th iteration. 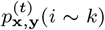 and 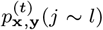 are the posterior coincidence probabilities for position pairs (*i, k*) and (*j, l*), respectively, from (*t*)th iteration. The extrinsic information 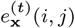 first sums all the base pair probabilities of alignable pairs from another one sequence with the co-incidence probabilities and then iterates over all the other sequences. *s***x**,**y** is the sequence identity for sequences **x** and **y**. The sequences with a low identity contribute more to the extrinsic information than sequences of higher identity. The sequence identity is defined as the fraction of nucleotides that are aligned and identical in the alignment.

### §6 LinearPartition for Base Pairing Probabilities Estimation with Extrinsic Information

The classical partition function algorithm scales cubically with sequence length. The slowness limits its extension to longer sequences. To address this bottleneck, our recent LinearPartition (46) algorithm approximates the partition function and base paring probability matrix computation in linear time. LinearPartition is significantly faster, and correlates better with the ground truth structures than the traditional cubic partition function calculation. Thus LinearTurboFold uses LinearPartition to predict base pair probabilities instead of the traditional *O*(*n*^3^)-time partition function.

TurboFold introduces the extrinsic information 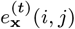 in the partition function as a pseudo-free energy term for each base pair (*i, j*). Similarly, in LinearPartition, for each span [*i, j*], which is the subsequence *x*_*i*_…*x*_*j*_, and its associated partition function *Q*(*i, j*), the partition function is modified as 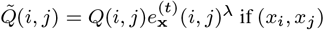 if (*x*_*i*_, *x*_*j*_) is an allowed pair, where *λ* denotes the contribution of the extrinsic information relative to the intrinsic information. Specifically, at each step *j*, among all possible spans [*i, j*] where *x*_*i*_ and *x*_*j*_ are paired, we replace the original partition function *Q*(*i, j*) with 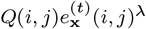 by multiplying the extrinsic information. Then LinearTurboFold applies the beam pruning heuristic over the modified partition function 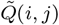 instead of the original.

Similarly, TurboFold II obtains the extrinsic information for all the *O*(*n*^2^) base pairs before the partition function calculation of each sequence, while only a linear number of base pairs survives in LinearPartition. Thus, LinearTurboFold only requires the extrinsic information for those promising base pairs that are visited in LinearPartition. Overall, for *k* homologous sequences, LinearTurboFold reduces the runtime of base pair probabilities estimation for each sequence from *O*(*kn*^3^ + *k*^2^*n*^2^) to 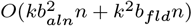 by applying the beam search *b*_*fld*_ to the partition function calculation, and only calculating extrinsic information for the saved base pairs.

### §7 MSA Generation and Secondary Structure Prediction

After several iterations, TurboFold II builds the multiple sequence alignment using a probabilistic consistency transformation, generating a guide tree and performing progressive alignment over the pairwise posterior co-incidence probabilities (30). The whole procedure is accelerated in virtue of the sparse matrix by discarding alignment pairs of probability smaller than a threshold (0.01 by default). Since LinearAlignment uses the beam search method and only saves a linear number of co-incident pairs, the MSA generation in LinearTurboFold costs linear runtime against the sequence length straightforwardly.

Estimated base pair probabilities are fed into downstream methods to predict secondary structures. To maintain the end-to-end linear-time property, LinearTurboFold uses ThreshKnot (49), which is a thresholded version of ProbKnot (68) and only considers base pairs of probability exceeding a threshold *θ* (*θ* = 0.3 by default). We evaluate the performance of ThreshKnot and MEA with different hyperparameters (*θ* and *γ*). On a sampled RNAStrAlign training set, ThreshKnot is closer to the upper right-hand than MEA, which indicates that ThreshKnot always has a higher Sensitivity than MEA at a given PPV (*SI Appendix*, Fig. S8B).

### §8 Efficiency and Scalability Datasets

Four datasets are built and used for measuring efficiency and scalability. To evaluate the efficiency and scalability of LinearTurboFold with sequence length, we collected groups of homologous RNA sequences with sequence length ranging from 200 *nt* to 29,903 *nt* with a fixed group size 5. Sequences are sampled from RNAStrAlign dataset (27), the Comparative RNA Web (CRW) Site (69), the Los Alamos HIV database (http://www.hiv.lanl.gov/) and the SARS-related betacoronaviruses (SARS-related) (53). RNAStrAlign, aggregated and released with TurboFold II, is an RNA alignment and structure database. Sequences in RNAStrAlign are categorized into families, i.e. sets of homologs, and some of families are further split into subfamilies. Each subfamily or family includes a multiple sequence alignment and ground truth structures for all the sequences. 20 groups of five homologs were randomly chosen from the small subunit ribosomal RNA (Alphaproteobacteria subfamily), SRP RNA (Protozoan subfamily), RNase P RNA (bacterial type A subfamily) and telomerase RNA families. For longer sequences, we sampled five groups of 23S rRNA (of sequence length ranging from 2,700 *nt* to 2,926 *nt*) from the CRW Site, HIV-1 genetic sequences (of sequence length ranging from 9,597 *nt* to 9,738 *nt*) from the Los Alamos HIV database, and SARS-related sequences (of sequence length ranging from 29,484 *nt* to 29,903 *nt*). All the sequences in one group belong to the same subfamily or subtype. We sampled five groups for each family and obtained 35 groups in total. Due to the runtime and memory limitations, we did not run TurboFold II on SARS-CoV-2 groups (Fig. 2, A and D).

To assess the runtime and memory usage of LinearTurboFold with group size, we fixed the sequence length around 1,500 *nt*, and sampled 5 groups of sequences from the small subunit ribosomal RNA (Alphaproteobacteria subfamily) with group size 5, 10, 15 and 20, respectively (Fig. 2, B and F). We used a Linux machine (CentOS 7.7.1908) with 2.30 GHz Intel Xeon E5-2695 v3 CPU and 755 GB memory, and gcc 4.8.5 for benchmarks.

We built a test set from the RNAStrAlign dataset to measure and compare the performance between LinearTurboFold and other methods. 60 groups of input sequences consisting of five homologous sequences were randomly selected from the small subunit ribosomal RNA (rRNA) (Alphaproteobacteria subfamily), SRP RNA (Protozoan subfamily), RNase P RNA (bacterial type A subfamily) and telomerase RNA families from RNAStrAlign dataset. We removed sequences shorter than 1,200 *nt* for the small subunit rRNA to filter out subdomains, and removed sequences that are shorter than 200 *nt* for SRP RNA following the TurboFold II paper to filter out less reliable sequences. We resampled the test set five times and show the average PPV, Sensitivity and F1 scores over the five samples (Fig. 2, C and F).

An RNAStrAlign training set was built to compare accuracies between MEA and ThreshKnot. 40 groups of 3, 5 and 7 homologs were randomly sampled from 5S ribosomal RNA (Eubacteria subfamily), group I intron (IC1 subfamily), tmRNA, and tRNA families from RNAStrAlign dataset. We chose *θ* = 0.1, 0.2, 0.3, 0.4 and 0.5 for ThreshKnot, and *γ* = 1, 1.5, 2, 2.5, 3, 3.5, 4, 8 and 16 for MEA. We reported the average secondary structure prediction accuracies (PPV and Sensitivity) across all training families (*SI Appendix*, Fig. S8B).

### §9 Benchmarks

The Sankoff algorithm (15) uses dynamic programming to simultaneously fold and align two or more sequences, and it requires *O*(*n*^3*k*^) time and *O*(*n*^2*k*^) space for *k* input sequences with the average length *n*. Both LocARNA (16) and MXSCARNA (18) are Sankoff-style algorithms.

LocARNA (local alignment of RNA) costs *O*(*n*^2^(*n*^2^ + *k*^2^)) time and *O*(*n*^2^ + *k*^2^) space by restricting the alignable regions. MXSCARNA progressively aligns multiple sequences as an extension of the pairwise alignment algorithm SCARNA (70) with improved score functions. SCARNA first aligns stem fragment candidates, then removes the inconsistent matching in the post-processing to generate the sequence alignment. MXSCARNA reduces runtime to *O*(*k*^3^*n*^2^) and space to *O*(*k*^2^*n*^2^) with a limited searching space of folding and alignment. Both MXSCARNA and LocARNA uses pre-computed base pair probabilities for each sequence as structural input. All the benchmarks use the default options and hyper-parameters running on the RNAStrAlign test set. TurboFold II iterates three times, then predicts secondary structures by MEA (*γ*=1). LinearTurboFold also runs three iterations with default beam sizes (*b*_*aln*_ = *b*_*fld*_ = 100) in LinearAlignment and LinearPartition, then predicts structures with ThreshKnot (*θ* = 0.3).

### §10 Significance Test

We use a paired, two-tailed permutation test (71) to measure the significant difference. Following the common practice, the repetition number is 10,000, and the significance threshold *α* is 0.05.

### §11 SARS-CoV-2 Datasets

We used two large SARS-CoV-2 datasets. The first dataset is used to draw a representative sample of most diverse SARS-CoV-2 genomes. We downloaded all the genomes submitted to GISAID (52) by December 29, 2020 (downloaded on December 29, 2020), and filtered out low-quality genomes (with more than 5% unknown characters and degenerate bases, shorter than 29,500 *nt*, or with framing error in the coding region), and we also discard genomes with more than 600 mutations compared with the SARS-CoV-2 reference sequence (NC_0405512.2) (72). After preprocessing, this dataset includes about 258,000 genomes. To identify a representative group of samples with more variable mutations, we designed a greedy algorithm to select 16 most diverse genomes genomes found at least twice in the 258,000 genomes. The general idea of the greedy algorithm is to choose genomes one by one with the most new mutations compared with the selected samples, which consists of only the reference sequence at the beginning.

The second, larger, dataset is to evaluate the conservation of regions with respect to more up-to-date variants. We did the same preprocessing as the first dataset on all the genomes submitted to GISAID by June 30, 2021 (downloaded on July 25, 2021). This resulted in a dataset of ∼2M genomes, which was used to evaluate conservation in Figure 5 and *SI Appendix*, Tab. S4–S6.

### §12 Data Availability

Our code, data and complete results for 25 SARS-CoV-2 and SARS-related genomes are released at: https://github.com/LinearFold/LinearTurboFold.

## ACKNOWLEDGMENTS

This work is supported in part by National Institutes of Health (R01 GM132185 to D.H.M.) and National Science Foundation (IIS-1817231 to L.H.).

## Footnotes

* Theoretically, the alignment part takes *O*(*k*^2^ *n*^2^) space. However, in practice, TurboFold II discards positions whose alignment co-incidence probabilities less than thresholds and only keeps a linear number of positions. (51)

† The average sequence identity is 0.9987 on that ∼2M dataset (downloaded on July 25, 2021).

