## Supplemental Material for "LinearTurboFold: Linear-Time Global Prediction of Conserved Structures for RNA Homologs with Applications to SARS-CoV-2"

PNAS
www.pnas.org

### Supplementary Information for

#### LinearTurboFold: Linear-Time Global Prediction of Conserved Structures for RNA Homologs with Applications to SARS-CoV-2

**Sizhen Li, He Zhang, Liang Zhang, Kai Zhao, Boxiang Liu, David H. Mathews*, and Liang Huang***

**Liang Huang:; David H. Mathews:**

##### This PDF file includes:

Figs. S1 to S14
Tables S1 to S7
Legend for Dataset S1
SI References

##### Other supplementary materials for this manuscript include the following:

Dataset S1

### Supporting Information

#### LinearTurboFold: Linear-Time Global Prediction of Conserved Structures for RNA Homologs with Applications to SARS-CoV-2

Sizhen Li, He Zhang, Liang Zhang, Kaibo Liu, Boxiang Liu, David H. Mathews, and Liang Huang

-

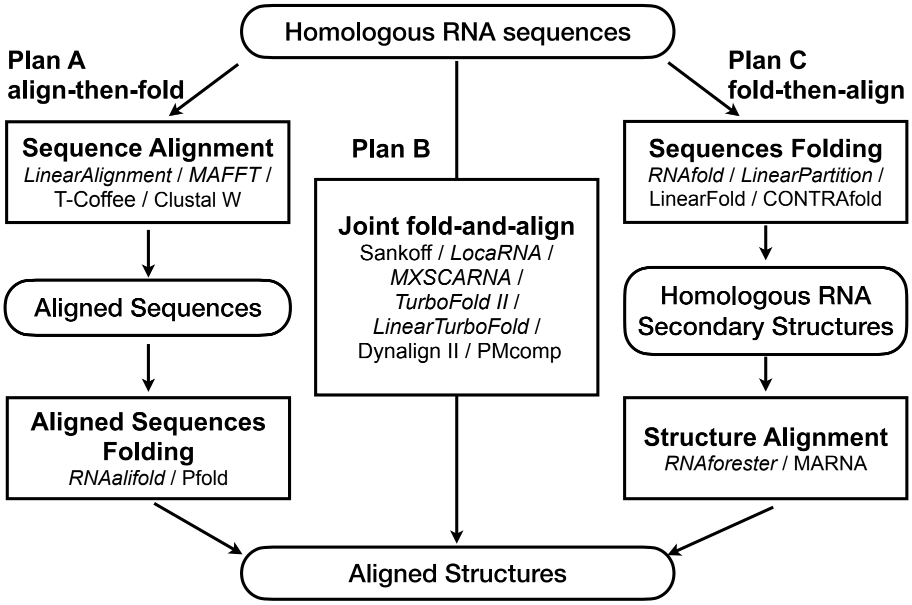


**Fig. S1.** Approaches of automate comparative sequence analysis can be categorized into three plans (1–3). Plan A, approach, involves two steps: first aligning sequences and then taking this alignment as input to predict the conserved structure. This was described by Waterman (4), and was subsequently refined and popularized by RNAalifold (5). This plan works well for homologs with a high sequence identity (Tab. S1). Plan C, "fold-then-align" approach, predicts plausible structures for the sequences first, and then aligns the structures to determine the sequence alignment and the optimal conserved structures. This was described by Waterman (6) and implemented in RNAforester (7) and MARNA (8). Italic methods in each plan are evaluated on RNAStrAlign dataset (Tab. S1). Plan B, "fold-and-align" approach, performs joint folding and alignment for multiple sequences, which requires more time and space. This was first proposed by Sankoff (9) using a dynamic programming algorithm. Several software packages provide implementations of the Sankoff algorithm (10–15) that use simplifications to reduce runtime.

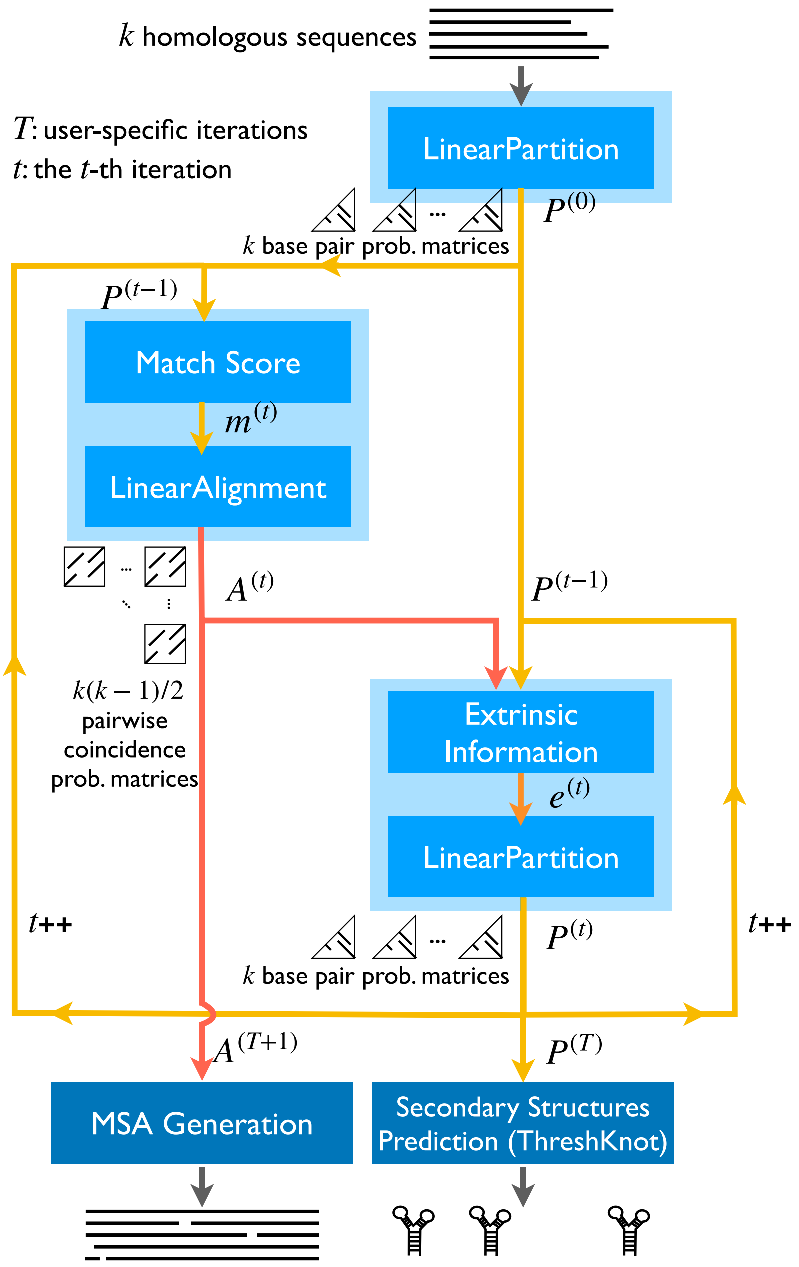


**Fig. S2.** The flowchart of LinearTurboFold with more detailed information. At the zeroth iteration, LinearPartition calculates the partition function and estimates the base pair probabilities for each sequence. From iteration 1 to $T$, the two major modules LinearAlignment and LinearPartition are conducted and updated in order with the match score and extrinsic information, respectively. The match score and extrinsic information are required and calculated for promising position pairs and base pairs during the LinearAlignment and LinearPartition computations, respectively. After $T$ iterations, the match score and LinearAlignment computations are performed one more time over the latest the base pair probabilities. A multiple sequence is generated based on the pairwise co-incidence probabilities from the ($T$+1)-th iteration, and secondary structures are predicted according to the base pair probabilities for each sequence from the $T$-th iteration.

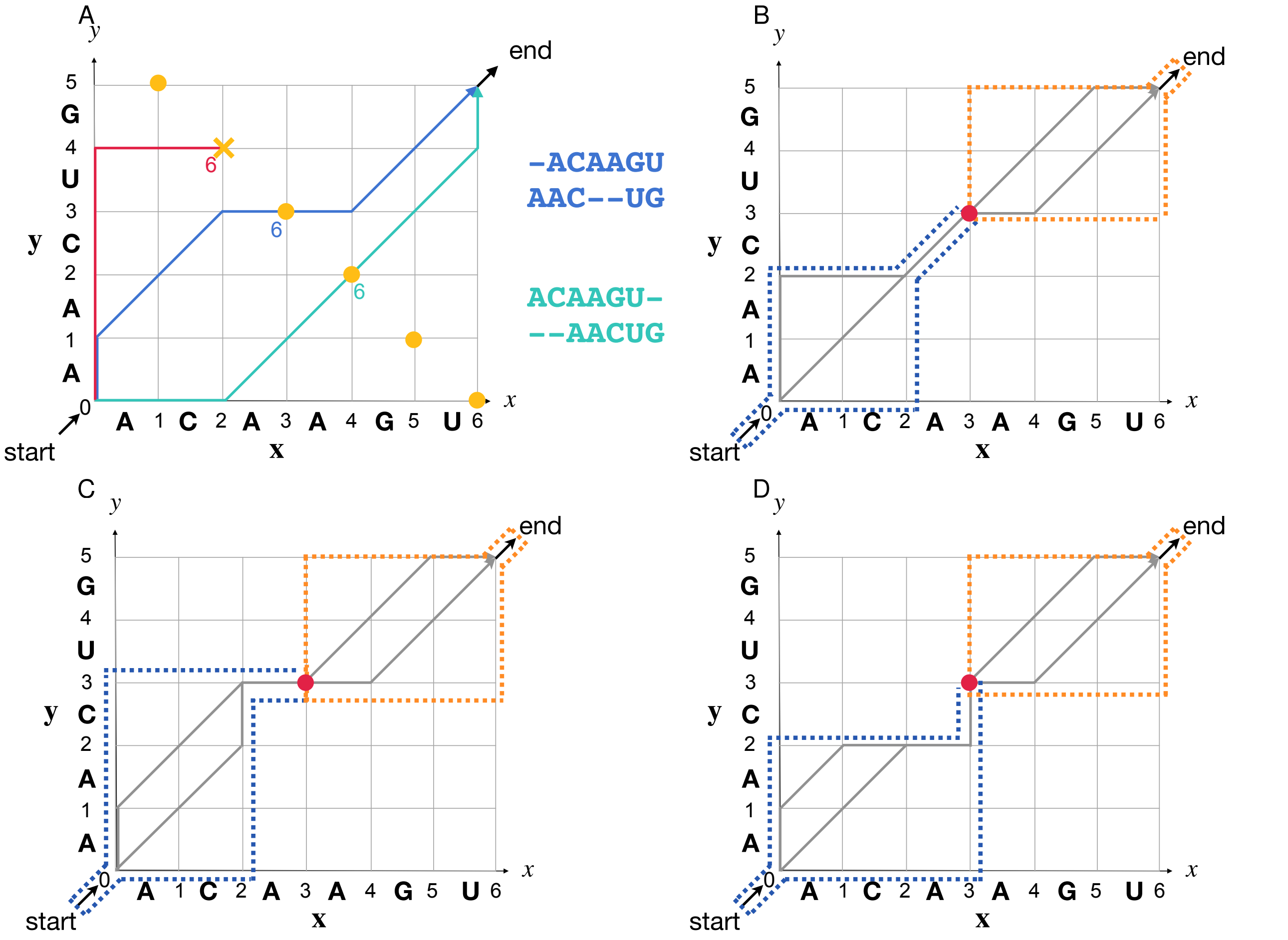


**Fig. S3.** Illustrations of LinearAlignment. $\mathbf{A}$: An example of aligning two sequences and the beam search method based on the step count. The $x$-axis and $y$-axis of the matrix represent two sequences $\mathbf{x}$ and $\mathbf{y}$. Yellow notes have the same step count 6. At step count 6, the red path is discarded because of its lower probability compared to others. There are two complete alignment paths (in green and blue) and the observed alignments are on the right side of the matrix with corresponding colors. $\mathbf{B}$: The area enclosed by the blue dashed line corresponds to $\alpha^{\text{ALN}}_{3,3}$, which includes all the partial alignments arriving at the node $(3,3)$ by the state $h$. And the region circled by the orange dashed line maintains all the partial alignments starting from the step $(\text{ALN}, 3, 3)$ ($\beta^{\text{ALN}}_{3,3}$). $\mathbf{C}$ and $\mathbf{D}$: The regions circled by the blue dashed lines are $\alpha^{\text{INS1}}_{3,3}$ and $\alpha^{\text{INS2}}_{3,3}$, and regions circled by the orange dashed lines are $\beta^{\text{INS1}}_{3,3}$ and $\beta^{\text{INS2}}_{3,3}$, respectively.

**function** FORWARD($\mathbf{x}$, $\mathbf{y}$, $b_{aln}$) ▷ the forward phase
$\alpha^{\text{ALN}}_{0,0} \leftarrow 1$ ▷ initial probability distribution
**for** $s = 0 ... |\mathbf{x}| + |\mathbf{y}| - 1$ **do** ▷ topological order
**for each** $h$ **in** $\mathcal{H}$ **do**
$B(s, h) \leftarrow$ all the nodes $(i, j)$ such that $\alpha^{h}_{i,j}$ exists and $i + j = s$
BEAMPRUNE($B(s, h), b_{aln}$) ▷ keep top $b_{aln}$ nodes in $B(s, h)$ by $\alpha^{h}_{i,j}$
**for each** node $(i, j)$ **in** $B(s, h)$ **do** ▷ transitions to next states
$\alpha^{\text{INS1}}_{i+1,j} \mathrel{+}= \alpha^{h}_{i,j} \cdot p_{\text{t}}(\text{INS1} \mid h) \cdot p_{\text{e}}((x_{i+1}, -) \mid \text{INS1})$
$\alpha^{\text{INS2}}_{i,j+1} \mathrel{+}= \alpha^{h}_{i,j} \cdot p_{\text{t}}(\text{INS2} \mid h) \cdot p_{\text{e}}((-, y_{j+1}) \mid \text{INS2})$
$\alpha^{\text{ALN}}_{i+1,j+1} \mathrel{+}= \alpha^{h}_{i,j} \cdot p_{\text{t}}(\text{ALN} \mid h) \cdot p_{\text{e}}((x_{i+1}, y_{j+1}) \mid \text{ALN})$
**return** $\alpha$

1. Pseudocode of the LinearAlignment algorithm forward phase

**function** BACKWARD($\mathbf{x}$, $\mathbf{y}$, $\alpha$, $B$) ▷ the backward phase
$\beta \leftarrow$ hash()
$\beta^{\text{ALN}}_{|\mathbf{x}|+1,|\mathbf{y}|+1} \leftarrow 1$ ▷ initialization
$p_{\mathbf{x},\mathbf{y}} \leftarrow \alpha^{\text{ALN}}_{|\mathbf{x}|+1,|\mathbf{y}|+1}$ ▷ probability of observing two sequences
$p_{i,j} \leftarrow 0$ ▷ co-incident probability initialization
**for** $s = |\mathbf{x}| + |\mathbf{y}| ... 0$ **do**
**for each** $h$ **in** $\mathcal{H}$ **do**
**for each** node $(i, j)$ **in** $B(s, h)$ **do** ▷ $B(s, h)$ saves $b_{aln}$ entries during the forward phase
**if** $i = |\mathbf{x}|$ **and** $j = |\mathbf{y}|$ **then** ▷ boundary conditions
$\beta^{h}_{i,j} = p_{\text{t}}(\text{ALN} \mid h) \cdot \beta^{\text{ALN}}_{|\mathbf{x}|+1,|\mathbf{y}|+1}$
**else**
$\beta^{h}_{i,j} = p_{\text{t}}(\text{ALN} \mid h) \cdot p_{\text{e}}((x_{i+1}, y_{j+1}) \mid \text{ALN}) \cdot \beta^{\text{ALN}}_{i+1,j+1}$
$+ p_{\text{t}}(\text{INS1} \mid h) \cdot p_{\text{e}}((x_{i+1}, -) \mid \text{INS1}) \cdot \beta^{\text{INS1}}_{i+1,j}$
$+ p_{\text{t}}(\text{INS2} \mid h) \cdot p_{\text{e}}((-, y_{j+1}) \mid \text{INS2}) \cdot \beta^{\text{INS2}}_{i,j+1}$
$p_{i,j} \mathrel{+}= \frac{\alpha^{h}_{i,j} \cdot \beta^{h}_{i,j}}{p_{\mathbf{x},\mathbf{y}}}$ ▷ update co-incident probabilities

2. Pseudocode of the LinearAlignment algorithm backward phase (co-incidence probability computation)

**Fig. S4.** The pseudocode of the LinearAlignment algorithm forward and backward phases (co-incidence probability computation). The pseudocode ignores boundary conditions for simplicity. LinearAlignment applies the beam search method first over $B(s, h)$, which is the collection of all the nodes $(i, j)$ with step count $s$ and the presence of $\alpha^{h}_{i,j}$ (the line in red). This algorithm only saves the top $b_{aln}$ nodes with the highest forward scores in $B(s, h)$, and these are subsequently allowed to make transitions to the next states. Here $b_{aln}$ is a user-specified beam size and the default value is 100. In total, $O(b_{aln}n)$ nodes survive because the length of $s$ is $|\mathbf{x}| + |\mathbf{y}|$ and each step count keeps $b_{aln}$ nodes.

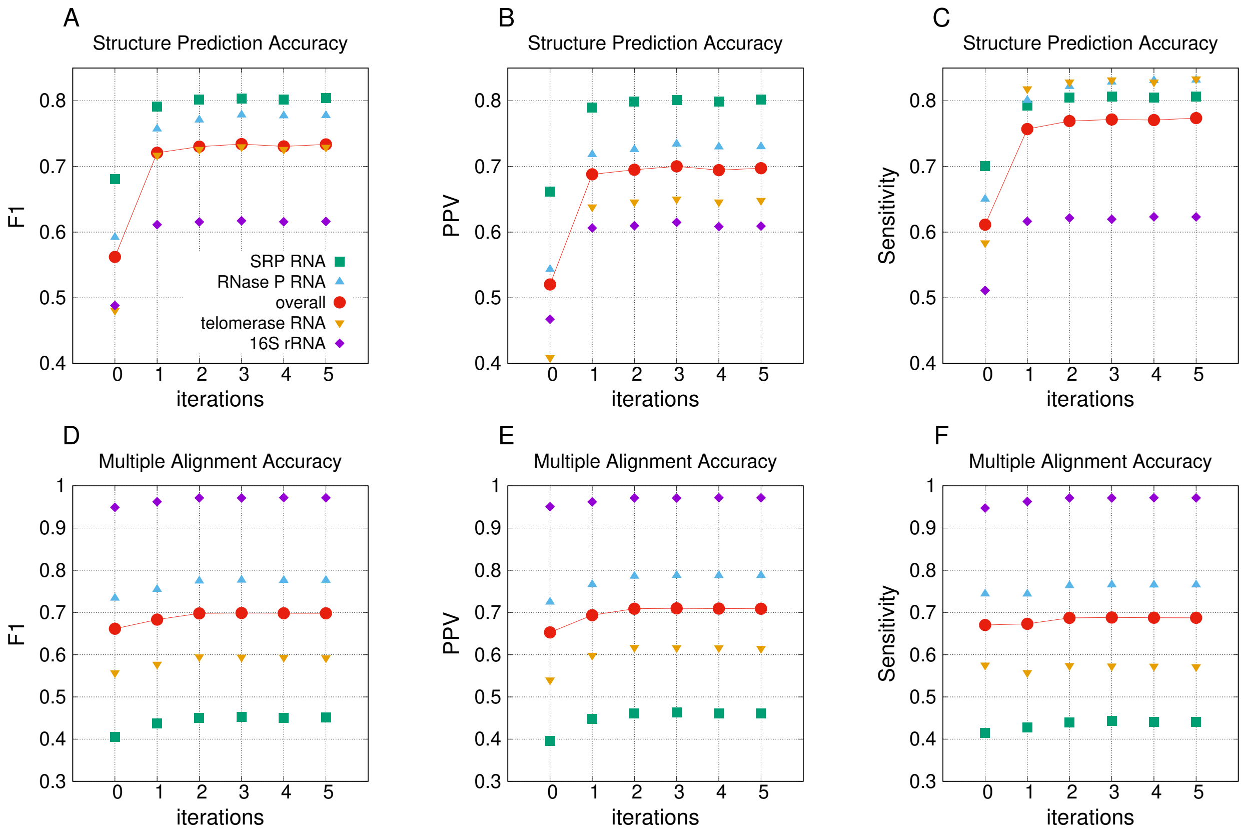


**Fig. S5.** Structure prediction and multiple sequence alignment accuracies against the number of iterations on the test set.

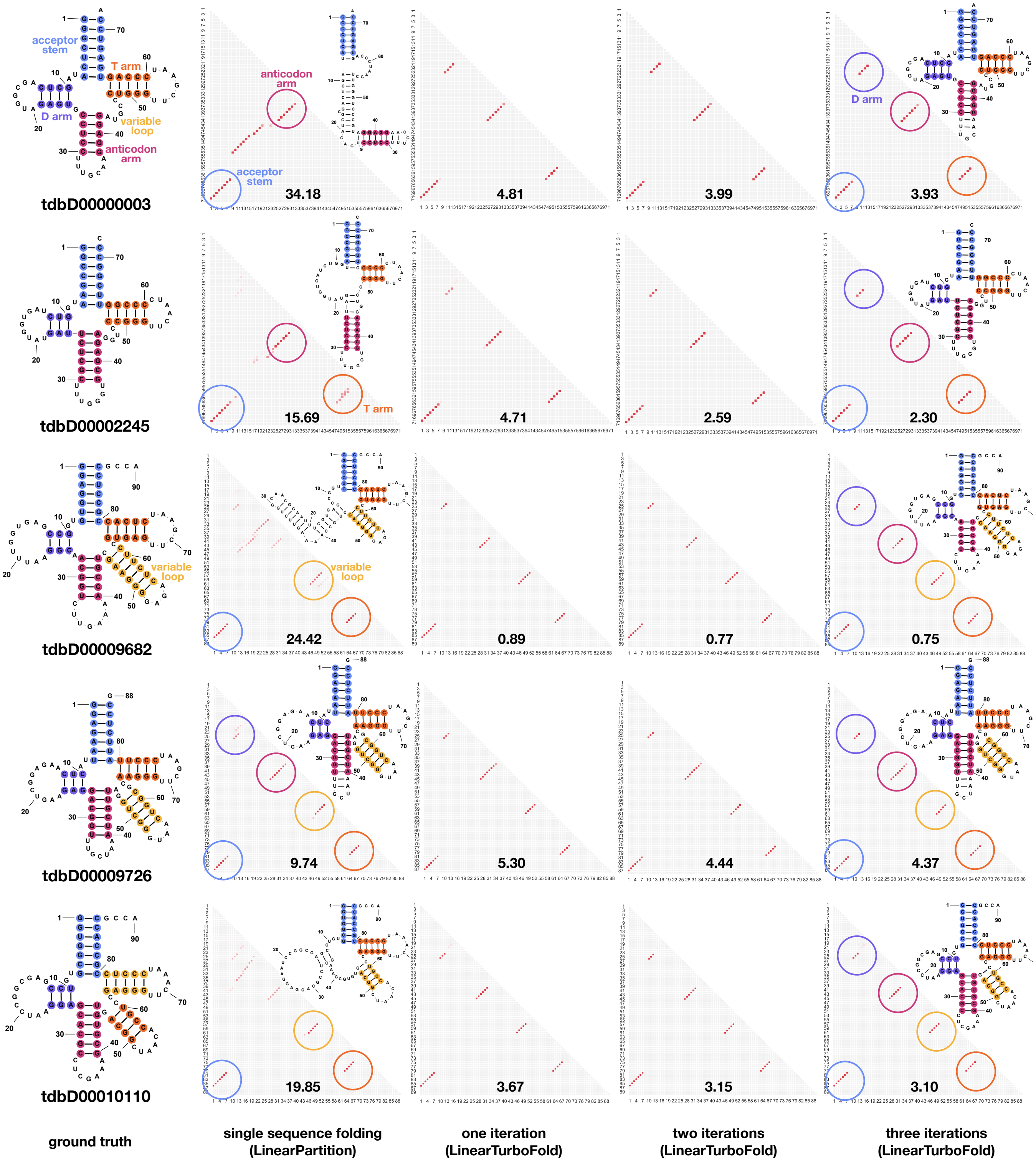


**Fig. S6.** Base pairing probabilities for a group of five tRNA sequences (tdbD00000003, tdbD00002245, tdbD00009682, tdbD00009726 and tdbD00010110). The first column shows the ground truth structures. The second column illustrates the base pair probabilities predicted by LinearPartition with corresponding ThreshKnot structures. The third to fifth columns show the base pair probabilities predicted by LinearTurboFold with different iterations. Ensemble defect values (16), which is the average number of incorrectly estimated paired nucleotides at equilibrium relative to the ground truth, are annotated on the bottom of lower triangles. We can observe that LinearPartition folds four sequences into wrong formations. By contrast, LinearTurboFold corrects the wrong predictions and sharpens the correct structures with larger probabilities iteratively, i.e., the ensemble defect values get smaller with more iterations. Alignment co-incidence probabilities between tdbD00002245 and tdbD00009726 across iterations are shown in Fig. S7.

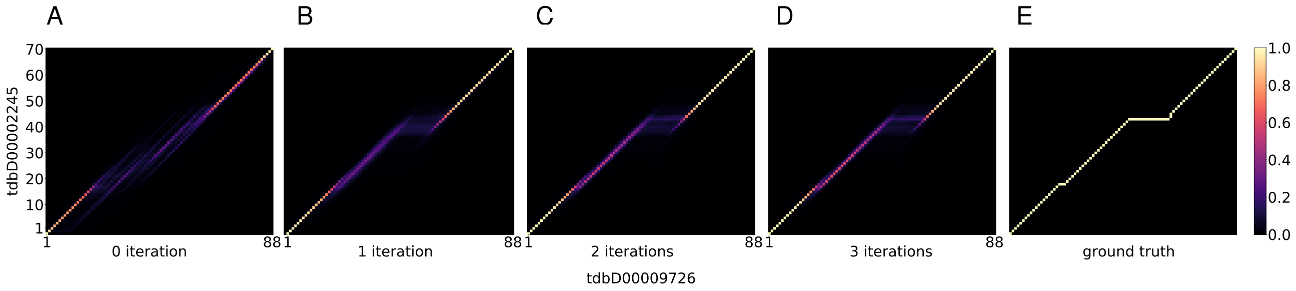


**Fig. S7.** Alignment co-incidence probabilities estimated by LinearTurboFold for a group of five tRNA sequences (tdbD00000003, tdbD00002245, tdbD00009726, tdbD00009682 and tdbD00010110) with different iterations. $\mathbf{A}$–$\mathbf{D}$: Alignment co-incidence probabilities between tdbD00002245 and tdbD00009726 from the zeroth iteration to the third iteration. The zeroth iteration means HMM sequence alignment without structural information. $\mathbf{E}$: The ground truth alignment. The variable loop is short and open in tdbD00002245, while a long stem loop inserted in this region in tdbD00009726 (see Fig. S6 for ground truth structures). As the iterations proceed, a long insertion is getting clearer around the variable loop in two sequences, and the probability matrix gets closer to the ground truth alignment.

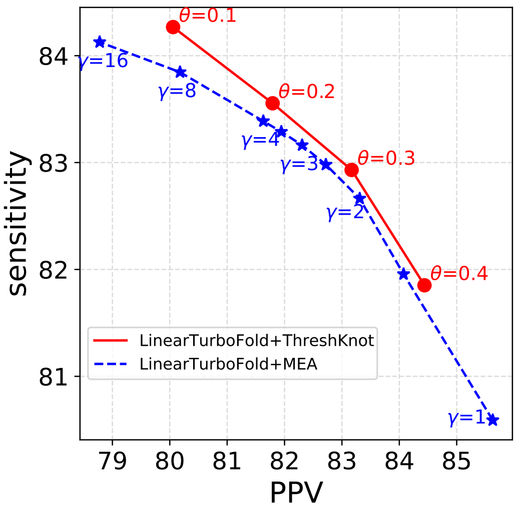


**Fig. S8.** Accuracy comparison between ThreshKnot and MEA on the training set with different hyper-parameters.

**Fig. S9.** Detailed information of the sampled 16 SARS-CoV-2 genomes and 9 SARS-related genomes. This dataset includes the reference sequences of SARS-CoV-2 and SARS-CoV-1 (NC_0405512.2, NC_004718.3). Most of the SARS-CoV-2 genomes include the D614G mutation, which has been a dominate mutation in the SARS-CoV-2 spike protein. B.1.1.7 lineage is a more infectious and lethal variant of SARS-CoV-2 first detected in the United Kingdom around November 2020. We utilized MAFFT (17) to generate the multiple sequence alignment and calculated the sequence identity with the reference sequence. The matrix under the table shows the pairwise sequence identities among 25 sequences.

| Accession ID | Species | Type | Submitted Date | Location | Length | Frequency | Mutations | Sequence identity | Note |
|---|---|---|---|---|---|---|---|---|---|
| NC_045512.2 | human | SARS-CoV-2 | 2020-01-17 | Wuhan, Asia | 29903 | 2 | - | - | - |
| EPI_ISL_454994 | human | SARS-CoV-2 | 2020-03-02 | Wuhan, Asia | 29864 | 3 | 36 | 0.999 | |
| EPI_ISL_572982 | human | SARS-CoV-2 | 2020-08-28 | England, Europe | 29882 | 2 | 28 | 0.999 | D614G |
| EPI_ISL_573173 | human | SARS-CoV-2 | 2020-09-08 | England, Europe | 29851 | 2 | 22 | 0.999 | D614G |
| EPI_ISL_573220 | human | SARS-CoV-2 | 2020-09-09 | England, Europe | 29784 | 2 | 19 | 0.999 | D614G |
| EPI_ISL_576666 | human | SARS-CoV-2 | 2020-09-18 | England, Europe | 29891 | 2 | 22 | 0.999 | D614G |
| EPI_ISL_639684 | human | SARS-CoV-2 | 2020-10-03 | Latvia, Europe | 29840 | 3 | 23 | 0.999 | D614G |
| EPI_ISL_648168 | human | SARS-CoV-2 | 2020-10-13 | Sweden, Europe | 29858 | 3 | 22 | 0.999 | D614G |
| EPI_ISL_706936 | human | SARS-CoV-2 | 2020-10-13 | England, Europe | 29828 | 2 | 23 | 0.999 | D614G |
| EPI_ISL_638950 | human | SARS-CoV-2 | 2020-10-14 | Scotland, Europe | 29891 | 2 | 29 | 0.999 | D614G |
| EPI_ISL_654499 | human | SARS-CoV-2 | 2020-10-20 | Sweden, Europe | 29876 | 3 | 20 | 0.999 | D614G |
| EPI_ISL_666966 | human | SARS-CoV-2 | 2020-10-30 | USA, NorthAmerica | 29879 | 2 | 23 | 0.999 | D614G |
| EPI_ISL_704698 | human | SARS-CoV-2 | 2020-11-01 | England, Europe | 29834 | 5 | 50 | 0.999 | D614G B.1.1.7 |
| EPI_ISL_723671 | human | SARS-CoV-2 | 2020-11-08 | England, Europe | 29876 | 2 | 32 | 0.999 | D614G |
| EPI_ISL_602304 | human | SARS-CoV-2 | 2020-11-12 | England, Europe | 29838 | 2 | 27 | 0.999 | D614G |
| EPI_ISL_710589 | human | SARS-CoV-2 | 2020-11-19 | Sweden, Europe | 29815 | 2 | 28 | 0.999 | D614G |
| NC_004718.3 | human | SARS-CoV-1 | 2003-04-13 | Vancouver, Canada | 29751 | - | 6277 | 0.789 | - |
| AY297028 | human | SARS-CoV-1 | 2003-05-19 | Beijing, Asia | 29715 | - | 6306 | 0.788 | - |
| AY515512.1 | human | SARS-CoV-1 | 2005-01-01 | Hong Kong, Asia | 29731 | - | 6298 | 0.788 | - |
| DQ182595.1 | human | SARS-CoV-1 | 2005-08-26 | Zhejiang, Asia | 29706 | - | 6298 | 0.788 | - |
| GU553363.1 | human | SARS-CoV-1 | 2010-01-15 | USA, NorthAmerica | 29644 | - | 6351 | 0.786 | - |
| DQ022305.2 | bat | SARS-CoV-1 | 2005-04-29 | Hong Kong, Asia | 29728 | - | 6337 | 0.787 | - |
| DQ648857.1 | bat | SARS-CoV-1 | 2006-05-23 | Hong Kong, Asia | 29741 | - | 6285 | 0.789 | - |
| EPI_ISL_402131 | bat | SARS-CoV-2 | 2013-07-24 | Yunnan, Asia | 29855 | - | 1176 | 0.961 | - |
| MG772934.1 | bat | SARS-CoV-1 | 2008-01-05 | Jiangsu, Asia | 29732 | - | 3740 | 0.874 | - |

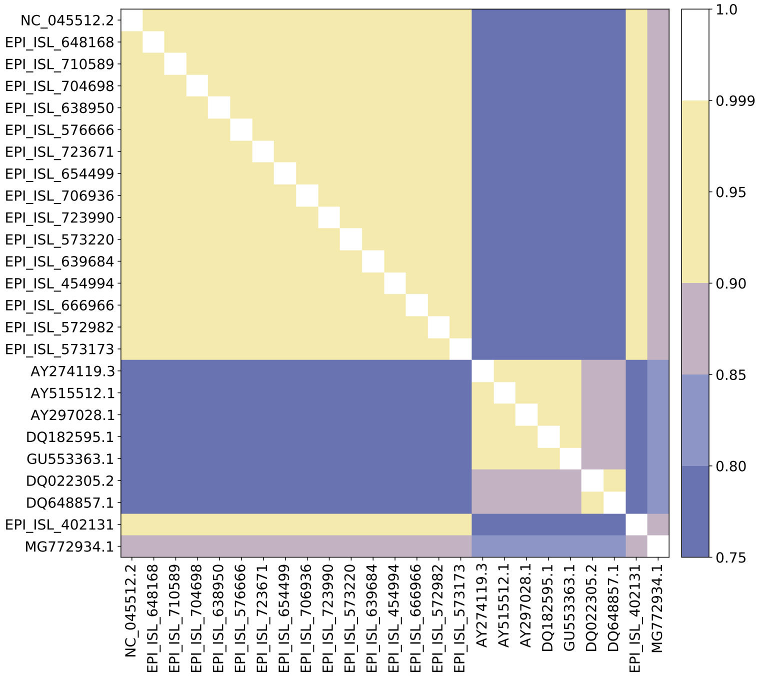


 Sizhen Li, He Zhang, Liang Zhang, Kai Zhao, Boxiang Liu, David H. Mathews*, and Liang Huang*

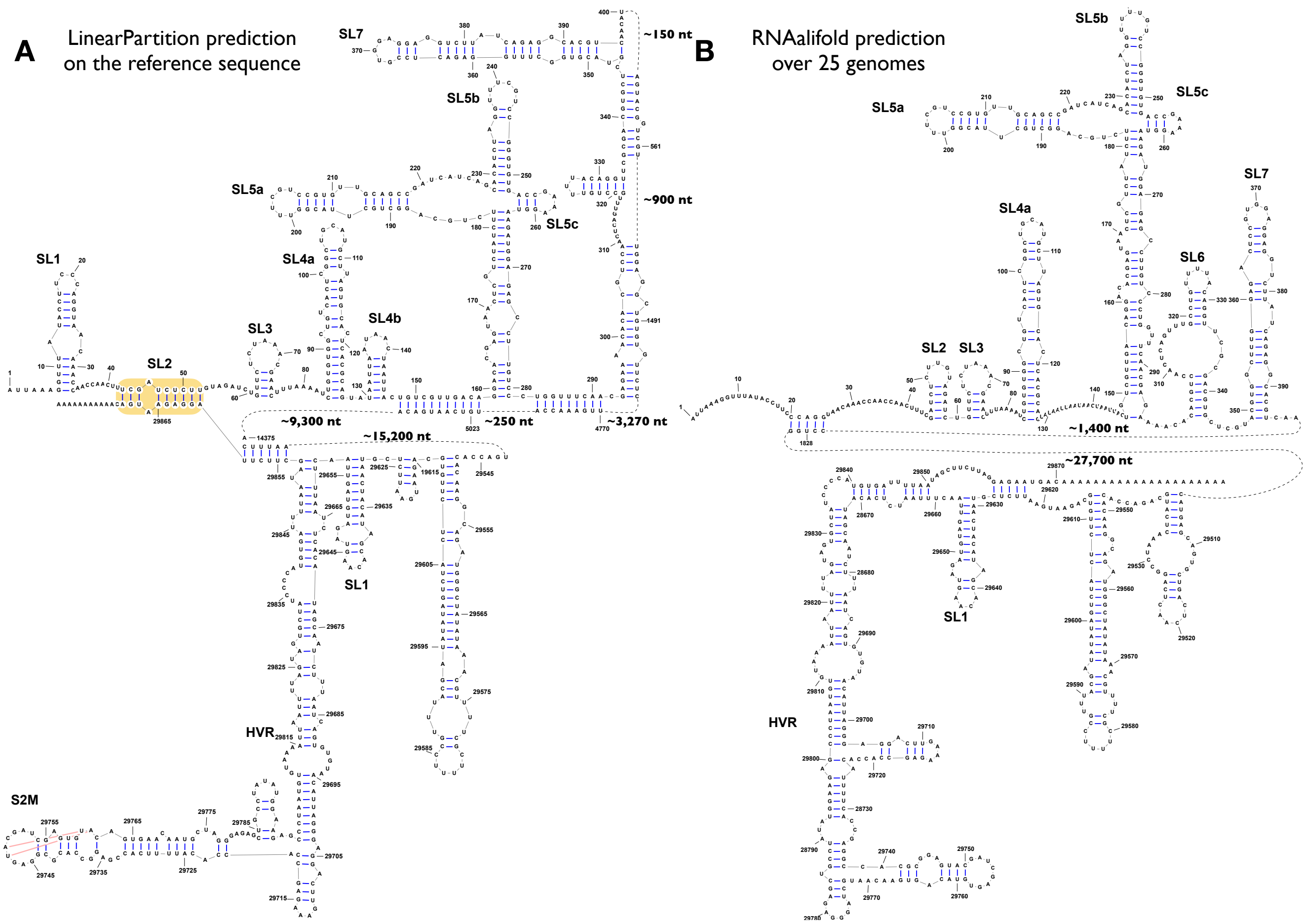


**Fig. S10.** Secondary structure prediction of SARS-CoV-2 for extended 5' and 3' UTRs. **A**: LinearPartition prediction of the SARS-CoV-2 reference sequence (NC_0405512.2) alone (single sequence folding). LinearPartition also predicts a long-range interaction between 5' and 3' UTRs. However, it involves the SL2 of the 5' UTR not SL3, which disagrees with LinearTurboFold prediction and Ziv *et al.*. **B**: RNAalifold (MFE) prediction for 25 genomes as listed in Fig. S9. RNAalifold did not find any 5'-3' pairs.

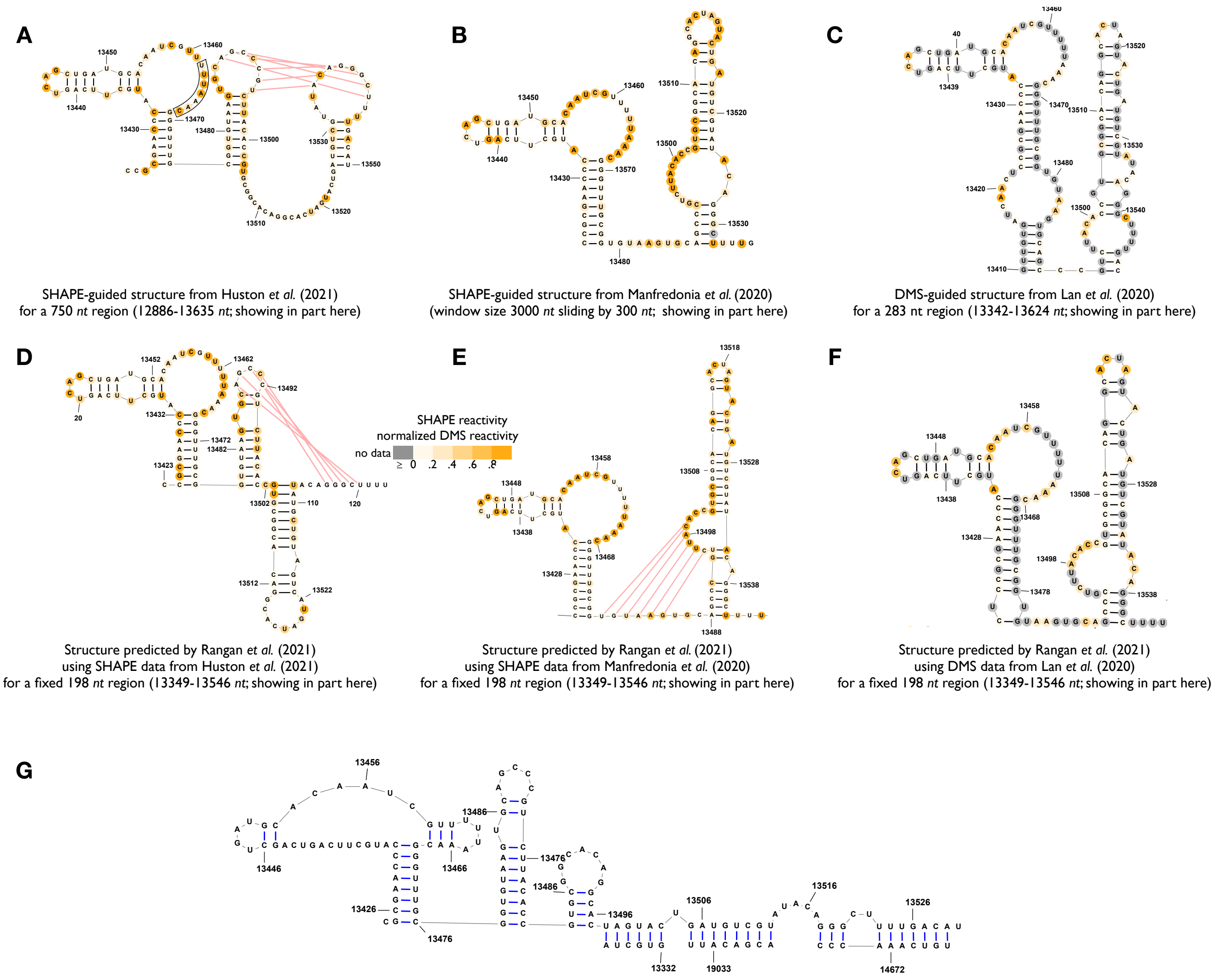


**Fig. S11.** Secondary structure predictions of SARS-CoV-2 for extended frameshifting stimulation element (13425-13545 $nt$). $\mathbf{A}$–$\mathbf{F}$: Experimentally-guided structures with different probing data for different regions. The structures in each column were estimated with the same experimental data but different regions. The structures in the second row were predicted by Rangan $et\ al.$ for a fixed region of 198 $nt$ (18) . G: RNAalifold (MFE) prediction over 25 genomes as listed in Fig. S9.

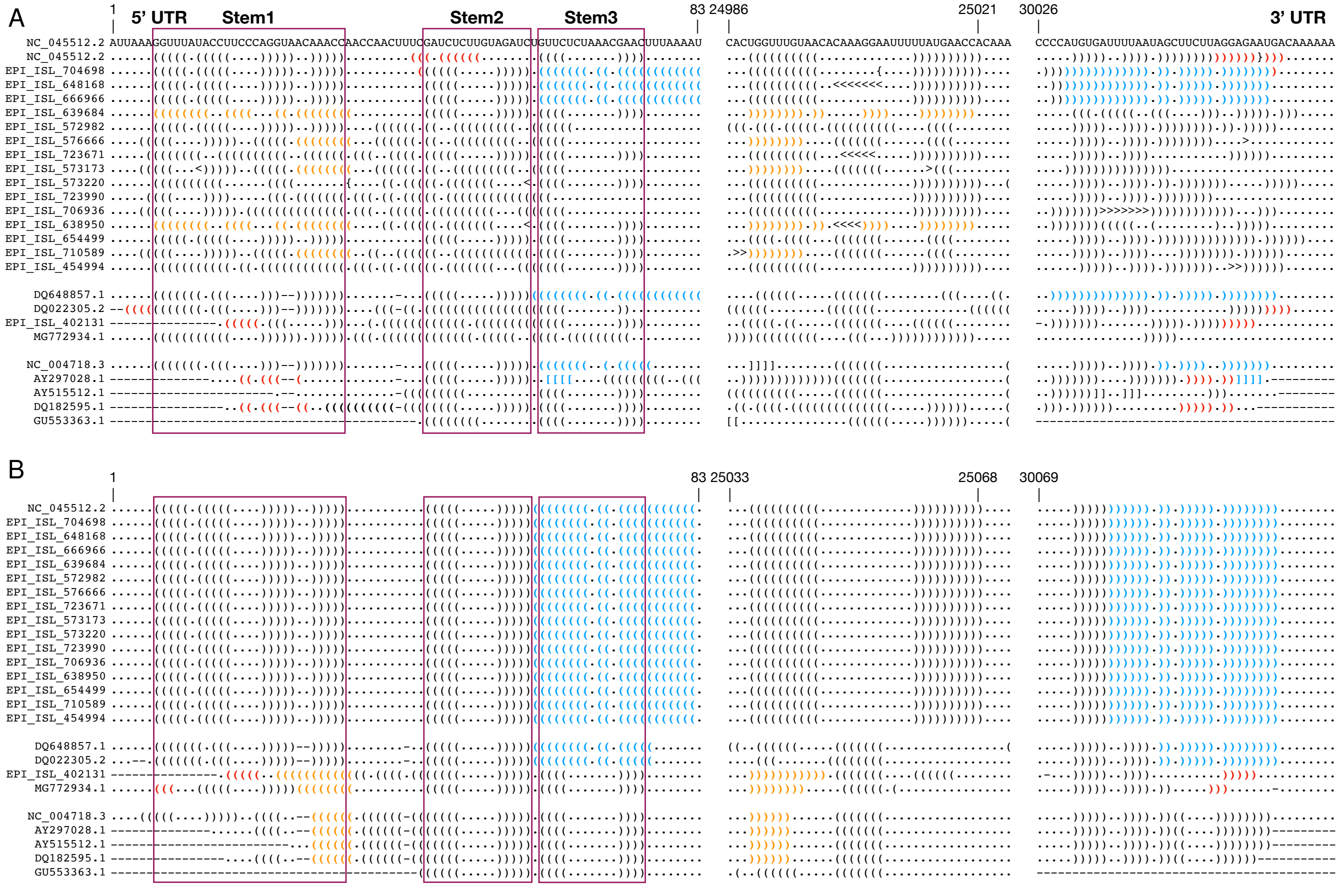


**Fig. S12.** ThreshKnot structures based on base-pairing probabilities from LinearPartition individually (**A**) and LinearTurboFold jointly (**B**), respectively, for 25 SARS-CoV-2 and SARS-related sequences as listed in Fig. S9. And these structures are aligned by the multiple sequence alignment from MAFFT (**A**) and LinearTurboFold (**B**), respectively. Three ranges (of ∼80 $nt$ at the start of 5'UTR, of ∼50 $nt$ at the end of 3' UTR and of ∼40 $nt$ around 25000 $nt$) are concatenated. The inter-actions between the first and third ranges are blue if they are the same as interactions detected by Ziv $et\ al.$ (19), otherwise, they are red. And the base pairs between the first and second ranges are orange. LinearPartitioncan model the same end-to-end interactions detected by Ziv $et\ al.$ for 4 out of 16 SARS-CoV-2 (EPI_ISL_704698, EPI_ISL_648168, EPI_ISL_666966 and EPI_ISL_639684) and 4 out of 9 SARS-related (DQ648857.1, NC_004718.3, AY297028.1 and DQ022305.2) sequences. The probabilities of such end-to-end pairs are relatively small for EPI_ISL_639684 and DQ022305.2 (Fig. S13), therefore the pairs are not shown in the threshKnot structures. For other sequences, however, LinearPartition either cannot predict the end-to-end interactions or it predicts them in wrong locations (e.g., EPI_ISL_572982 and NC_045512.2). By contrast, LinearTurboFold propagates the correct structural information, i.e., base-pairing probabilities (Fig. S13), from these eight sequences to other homologs, and finally folds all the SARS-CoV-2 and two SARS-related sequences into the same structure. Meanwhile, LinearTurboFold predicts a long-range interaction involving the SL1 of 5' UTR and a segment around 25000 $nt$ for other six SARS-related seqmuences. One SARS-relatedsequence (GU553363.1) can form neither interactions due to incomplete UTRs.

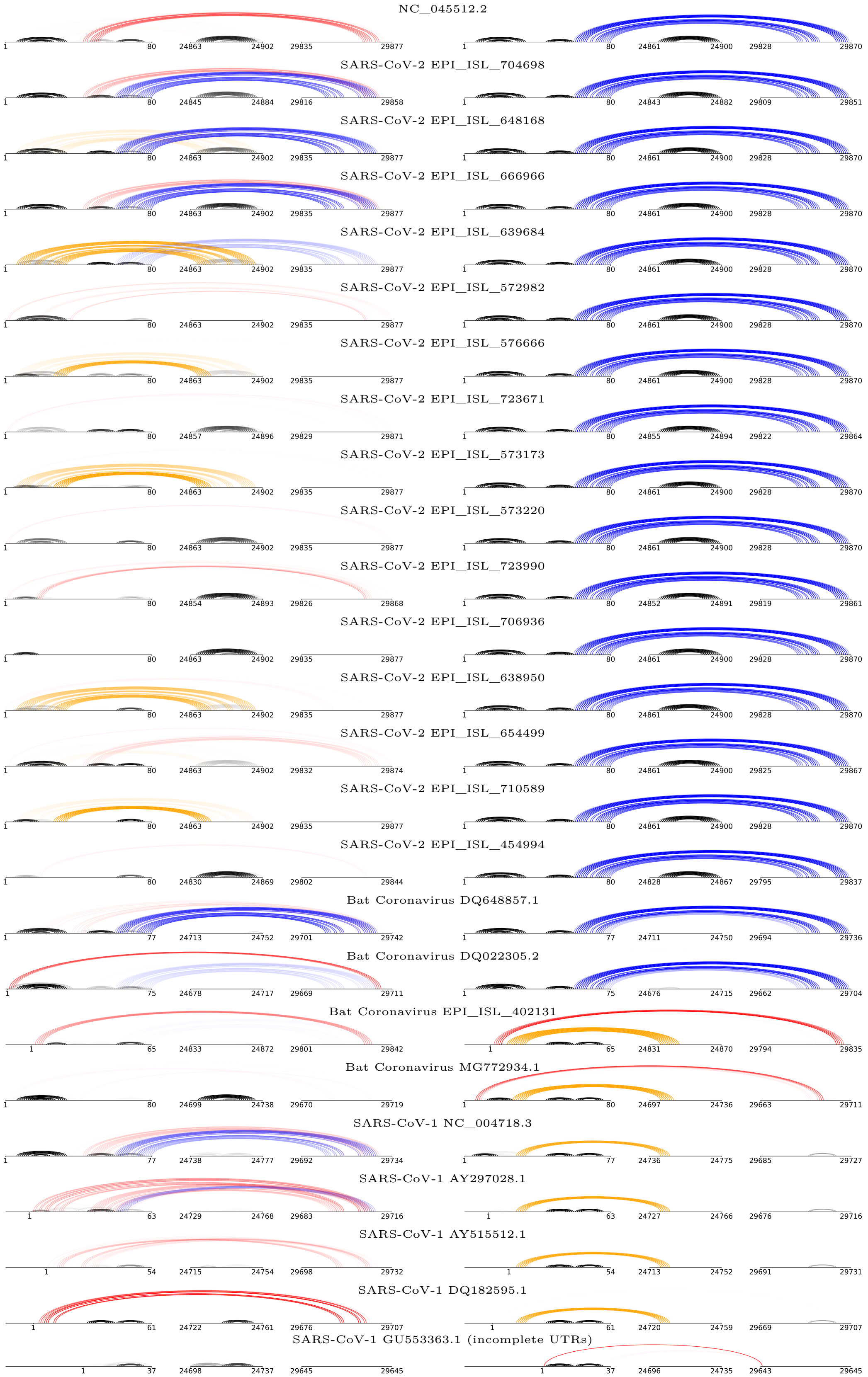


**Fig. S13.** Base-pairing probabilities predicted by LinearPartition individually (left column) and LinearTurboFold jointly (right column), respectively, for 25 SARS-CoV-2 and SARS-related genomes. The same three ranges as shown in Fig. S12 are concatenated. The arcs represent base pairs, whose transparency shows corresponding probabilities.

```
>NC_045512.2_Wuhan_seafood_market_pneumonia_virus_isolate_Wuhan-Hu-1__complete_genome:
CGUUCGGAUGCUCGAACUGCACCUCAUGGUCAUGUUAUGGUUGAGCUGGUAGCAGAACUCGAAGGCAUUCAGUACGGUCGUAGUGGUGAGACACUUGGUGUCCUUGUCCCUCAUGUGGGCGAAAUACCAGUGGCUUACCGC
(((.(((((((((((.((((((((((...((((...)))).)))...))).))))...)))..))))))).).)))......((((((((.((((.(((((..(((.(((......)))))).))))))))).))))))))
>hCoV-19_Sweden_20-53189_2020_EPI_ISL_648168_2020-10-13_Europe:
CGUUCGGAUGCUCGAACUGCACCUCAUGGUCAUGUUAUGGUUGAGCUGGUAGCAGAACUCGAAGGCAUUCAGUACGGUCGUAGUGGUGAGACACUUGGUGUCCUUGUCCCUCAUGUGGGCGAAAUACCAGUGGCUUACCGC
(((.(((((((((((.((((((((((...((((...)))).)))...))).))))...)))..))))))).).)))......((((((((.((((.(((((..(((.(((......)))))).))))))))).))))))))
>hCoV-19_Sweden_20-53470_2020_EPI_ISL_710589_2020-11-19_Europe:
CGUUCGGAUGCUCGAACUGCACCUCAUGGUCAUGUUAUGGUUGAGCUGGUAGCAGAACUCGAAGGCAUUCAGUACGGUCGUAGUGGUGAGACACUUGGUGUCCUUGUCCCUCAUGUGGGCGAAAUACCAGUGGCUUACCGC
(((.(((((((((((.((((((((((...((((...)))).)))...))).))))...)))..))))))).).)))......((((((((.((((.(((((..(((.(((......)))))).))))))))).))))))))
>hCoV-19_England_MILK-B3D791_2020_EPI_ISL_704698_2020-11-01_Europe:
CGUUCGGAUGCUCGAACUGCACCUCAUGGUCAUGUUAUGGUUGAGCUGGUAGCAGAACUCGAAGGCAUUCAGUACGGUCGUAGUGGUGAGACACUUGGUGUCCUUGUCCCUCAUGUGGGCGAAAUACCAGUGGCUUACCGC
(((.(((((((((((.((((((((((...((((...)))).)))...))).))))...)))..))))))).).)))......((((((((.((((.(((((..(((.(((......)))))).))))))))).))))))))
>hCoV-19_Scotland_QEUH-A77C42_2020_EPI_ISL_638950_2020-10-14_Europe:
CGUUCGGAUGCUCGAACUGCACCUCAUGGUCAUGUUAUGGUUGAGCUGGUAGCAGAACUCGAAGGCAUUCAGUACGGUCGUAGUGGUGAGACACUUGGUGUCCUUGUCCCUCAUGUGGGCGAAAUACCAGUGGCUUACCGC
(((.(((((((((((.((((((((((...((((...)))).)))...))).))))...)))..))))))).).)))......((((((((.((((.(((((..((((.((......)))))).))))))))).))))))))
>hCoV-19_England_ALDP-9D8D3E_2020_EPI_ISL_576666_2020-09-18_Europe:
CGUUCGGAUGCUCGAACUGCACCUCAUGGUCAUGUUAUGGUUGAGCUGGUAGCAGAACUCGAAGGCAUUCAGUACGGUCGUAGUGGUGAGACACUUGGUGUCCUUGUCCCUCAUGUGGGCGAAAUACCAGUGGCUUACCGC
(((.(((((((((((.((((((((((...((((...)))).)))...))).))))...)))..))))))).).)))......((((((((.((((.(((((..((((.((......)))))).))))))))).))))))))
>hCoV-19_England_ALDP-B5E5FA_2020_EPI_ISL_723671_2020-11-08_Europe:
CGUUCGGAUGCUCGAACUGCACCUCAUGGUCAUGUUAUGGUUGAGCUGGUAGCAGAACUCGAAGGCAUUCAGUACGGUCGUAGUGGUGAGACACUUGGUGUCCUUGUCCCUCAUGUGGGCGAAAUACCAGUGGCUUACCGC
(((.(((((((((((.((((((((((...((((...)))).)))...))).))))...)))..))))))).).)))......((((((((.((((.(((((..(((((.(......)))))).))))))))).))))))))
>hCoV-19_Sweden_20-09856_2020_EPI_ISL_654499_2020-10-20_Europe:
CGUUCGGAUGCUCGAACUGCACCUCAUGGUCAUGUUAUGGUUGAGCUGGUAGCAGAACUCGAAGGCAUUCAGUACGGUCGUAGUGGUGAGACACUUGGUGUCCUUGUCCCUCAUGUGGGCGAAAUACCAGUGGCUUACCGC
(((.(((((((((((.((((((((((...((((...)))).)))...))).))))...)))..))))))).).)))......((((((((.((((.(((((..(((.(((......)))))).))))))))).))))))))
>hCoV-19_England_QEUH-A61A50_2020_EPI_ISL_706936_2020-10-13_Europe:
CGUUCGGAUGCUCGAACUGCACCUCAUGGUCAUGUUAUGGUUGAGCUGGUAGCAGAACUCGAAGGCAUUCAGUACGGUCGUAGUGGUGAGACACUUGGUGUCCUUGUCCCUCAUGUGGGCGAAAUACCAGUGGCUUACCGC
(((.(((((((((((.((((((((((...(((.....))).)))...))).))))...)))..))))))).).)))......((((((((.((((.(((((..(((.(((......)))))).))))))))).))))))))
>hCoV-19_England_CAMC-B7B1B7_2020_EPI_ISL_723990_2020-11-12_Europe:
CGUUCGGAUGCUCGAACUGCACCUCAUGGUCAUGUUAUGGUUGAGCUGGUAGCAGAACUCGAAGGCAUUCAGUACGGUCGUAGUGGUGAGACACUUGGUGUCCUUGUCCCUCAUGUGGGCGAAAUACCAGUGGCUUACCGC
(((.(((((((((((.((((((((((...((((...)))).)))...))).))))...)))..))))))).).)))......((((((((.((((.(((((..(((.(((......)))))).))))))))).))))))))
>hCoV-19_England_CAMC-9C1B01_2020_EPI_ISL_573220_2020-09-09_Europe:
CGUUCGGAUGCUCGAACUGCACCUCAUGGUCAUGUUAUGGUUGAGCUGGUAGCAGAACUCGAAGGCAUUCAGUACGGUCGUAGUGGUGAGACACUUGGUGUCCUUGUCCCUCAUGUGGGCGAAAUACCAGUGGCUUACCGU
(((.(((((((((((.((((((((((...((((...)))).)))...))).))))...)))..))))))).).))).......(((((((.((((.(((((..(((.(((......)))))).))))))))).))))))).
>hCoV-19_Latvia_187_2020_EPI_ISL_639684_2020-10-03_Europe:
CGUUCGGAUGCUCGAACUGCACCUCAUGGUCAUGUUAUGGUUGAGCUGGUAGCAGAACUCGAAGGCAUUCAGUACGGUCGUAGUGGUGAGACACUUGGUGUCCUUGUCCCUCAUGUGGGCGAAAUACCAGUGGCUUACCGC
(((.(((((((((((.((((((((((...((((...)))).)))...))).))))...)))..))))))).).)))......((((((((.((((.(((((..((((((.......)))))).))))))))).))))))))
>hCoV-19_Wuhan_HB-WHCM-114_2020_EPI_ISL_454994_2020-03-02_Asia:
CGUUCGGAUGCUCGAACU---------------------------------GCAGAACUCGAAGGCAUUCAGUACGGUCGUAGUGGUGAGACACUUGGUGUCCUUGUCCCUCAUGUGGGCGAAAUACCAGUGGCUUACCGC
(((.(((((((((((...---------------------------------.......)))..))))))).).)))......((((((((.((((.(((((..(((.(((......)))))).))))))))).))))))))
>hCoV-19_USA_CA-ALSR-4490_2020_EPI_ISL_666966_2020-10-30_NorUhAmerica:
CGUUCGGAUGCUCGAACUGCACCUCAUGGUCAUGUUAUGGUUGAGCUGGUAGCAGAACUCGAAGGCAUUCAGUACGGUCGUAGUGGUGAGACACUUGGUGUCCUUGUCCCUCAUGUGGGCGAAAUACCAGUGGCUUACCGC
(((.(((((((((((.((((((((((...((((...)))).)))...))).))))...)))..))))))).).)))......((((((((.((((.(((((..((((((.......)))))).))))))))).))))))))
>hCoV-19_England_ALDP-9B601F_2020_EPI_ISL_572982_2020-08-28_Europe:
CGUUCGGAUGCUCGAACUGCACCUCAUGGUCAUGUUAUGGUUGAGCUGGUAGCAGAACUCGAAGGCAUUCAGUACGGUCGUAGUGGUGAGACACUUGGUGUCCUUGUCCCUCAUGUGGGCGAAAUACCAGUGGCUUACCGC
(((.(((((((((((.((((((((((...((((...)))).)))...))).))))...)))..))))))).).)))......((((((((.((((.(((((..(((.(((......)))))).))))))))).))))))))
>hCoV-19_England_CAMC-9C01A6_2020_EPI_ISL_573173_2020-09-08_Europe:
CGUUCGGAUGCUCGAACUGCACCUCAUGGUCAUGUUAUGGUUGAGCUGGUAGCAGAACUCGAAGGCAUUCAGUACGGUCGUAGUGGUGAGACACUUGGUGUCCUUGUCCCUCAUGUGGGCGAAAUACCAGUGGCUUACCGC
(((.(((((((((((.((((((((((...((((...)))).)))...))).))))...)))..))))))).).)))......((((((((.((((.(((((..(((.(((......)))))).))))))))).))))))))
>NC_004718.3_SARS_coronavirus_Tor2__complete genome:
CGUUCUGAUGCCUUAAGCACCAAUCACGGCCACAAGGUCGUUGAGCUGGUUGCAGAAAUGGACGGCAUUCAGUACGGUCGUAGCGGUAUAACACUGGGAGUACUCGUGCCACAUGUGGGCGAAACCCCAAUUGCAUACCGC
(((.((((((((.(..((((((.((((((((....))))).)))..)))).))......)...))))).))).)))......(((((((.....((((.((..((((.(((....))))))).)))))).....)))))))
>GU553363.1_SARS_coronavirus_HKU-39849_isolate_TCVSP-HARROD-00001__complete_genome:
CGUUCUGAUGCCUUAAGCACCAAUCACGGCCACAAGGUCGUUGAGCUGGUUGCAGAAAUGGACGGCAUUCAGUACGGUCGUAGCGGUAUAACACUGGGAGUACUCGUGCCACAUGUGGGCGAAACCCCAAUUGCAUACCGC
(((.((((((((.(..((((((.((((((((....))))).)))..)))).))......)...))))).))).)))......(((((((.....((((.((..((((.(((....))))))).)))))).....)))))))
>AY515512.1_SARS_coronavirus_HC_SZ_61_03__complete_genome:
CGUUCUGAUGCCUUAAGCACCAAUCACGGCCACAAGGUCGUUGAGCUGGUUGCAGAAAUGGACGGCAUUCAGUACGGUCGUAGCGGUAUAACACUGGGAGUACUCGUGCCACAUGUGGGCGAAACCCCAAUUGCAUACCGC
(((.((((((((.((.((((((.((((((((....))))).)))..)))).)).....))...))))).))).)))......(((((((.....((((.((..((((.(((....))))))).)))))).....)))))))
>AY297028.1_SARS_coronavirus_ZJ01__complete_genome:
CGUUCUGAUGCCUAAAGCACCUAUCACGGCCACAAGGUCGUUGAGCUGGUUGCAGAAAUGGACGGCAUUCAGUACGGUCGUAGCGGUAUAACACUGGGAGUACUCGUGCCACAUGUGGGCGAAACCCCAAUUGCAUACCGC
(((.((((((((....(((((..((((((((....))))).)))...))).))..........))))).))).)))......(((((((.....((((.((..((((.(((....))))))).)))))).....)))))))
>DQ182595.1_SARS_coronavirus_ZJ0301_from_China__complete_genome:
CGUUCUGAUGCCUUAAGCACCAAUCACGGCCACAAGGUCGUUGAGCUGGUUGCAGAAAUGGACGGCAUUCAGUACGGUCGUAGCGGUAUAACACUGGGAGUACUCGUGCCACAUGUGGGCGAAACCCCAAUUGCAUACCGC
(((.((((((((.((.((((((.((((((((....))))).)))..)))).)).....))...))))).))).)))......(((((((.....((((.((..((((.(((....))))))).)))))).....)))))))
>MG772934.1_Bat_SARS-like_coronavirus_isolate_bat-SL-CoVZXC21__complete_genome:
CGUUCUGAUGCCCGAACUGCACAUCAUGGCCAUGUUAUGGUCGAAUUAGUAGCAGAACUCGAUGGCAUUCAGUACGGUCGUAGUGGUGAGACACUUGGUGUUCUUGUCCCUCAUGUGGGAGAGGUACCAGUUGCUUACCGU
(((.(((((((((((.((((((....(((((((...)))))))....)).))))...)))..))))).))).))).......(((((((..(((.((((..(((.((((......)))).))))))))))..))))))).
>DQ022305.2_Bat_SARS_coronavirus_HKU3-1__complete_genome:
CGAUCUGAUGCUCUAAGCACCAAUCACGGCCACAAGGUUGUGGAAUUGGUUGCAGAAUUAGAUGGCAUUCAGUUCGGUCGUAGCGGUAUAACACUGGGAGUUCUCGUGCCACAUGUGGGCGAAACCCCAAUCGCAUACCGU
(((.(((((((((((((((((((((((((((....)))))))..)))))).))....)))).))))).))).)))......(((((((.....((((.(((.((((.(((....)))))))))))))).....)))))))
>DQ648857.1_Bat_coronavirus_(BtCoV_279_2005)__complete_genome:
CGUUCUGAUGCCCAGGGCACCAAUCACGGCUAUAAGGUCGUUGAAUUGGUUGCAGAAUUGGACGGCAUUCAGUACGGUCGUAGCGGUACAACACUGGGAGUCCUCGUGCCACAUGUGGGCGAAACCCCAGUUGCGUACCGU
(((.((((((((..(((((((((((((((((....)))))).)).))))).)).....))...))))).))).)))......(((((((...((((((.((..((((.(((....))))))).))))))))...)))))))
>BetaCoV_bat_Yunnan_RatG13_2013_EPI_ISL_402131:
CGUUCUGAUGCUCGAACUGCACCUCAUGGCCAUGUUAUGGUUGAGCUGGUAGCAGAACUUAAUGGCAUUCAGUAUGGUCGUAGUGGUGAGACACUCGGUGUCCUUGUCCCUUAUGUGGGCGAAACACCAGUGGUUUACCGC
(((.((((((((.((.(((((((((..((((((...))))))))...))).))))...))...))))).))).)))......((((((((.((((.(((((..(((((((....).)))))).))))))))).))))))))
```

**Fig. S14.** A glimpse of the whole MSA and aligned predicted structures for 25 genomes from LinearTurboFold, which are available in https://github.com/LinearFold/LinearTurboFold/blob/main/sars-cov-2_results/. Each genome corresponds to three lines: name, aligned sequence and aligned structure, respectively.

**Table S1. Structure prediction and multiple sequence alignment accuracies. LinearTurboFold (1) and (2) use the same beam size (100) in LinearPartition, and different beam sizes of 100 and $\infty$ (i.e., exact search) in HMM alignment, respectively.**

| | **Structure Prediction Accuracy** | | | | | | | |
|---|---|---|---|---|---|---|---|---|
| | LinearTurboFold (1) | (2) | TurboFold II | LocARNA | MXSCARNA | LinearPartition | Vienna RNAfold | RNAalifold |
| | | | | | PPV | | | |
| SRP | 0.801 | 0.801 | **0.819** | 0.698 | 0.485 | 0.662 | 0.673 | 0.629 |
| telomerase | 0.650 | 0.649 | **0.685** | 0.516 | 0.465 | 0.409 | 0.430 | 0.602 |
| RNase P RNA | 0.734 | 0.731 | **0.752** | 0.606 | 0.573 | 0.543 | 0.571 | 0.698 |
| 16S rRNA | 0.615 | 0.610 | 0.608 | 0.586 | **0.662** | 0.467 | 0.464 | 0.628 |
| overall | 0.700 | 0.698 | **0.716** | 0.602 | 0.546 | 0.520 | 0.534 | 0.639 |
| | | | | | Sensitivity | | | |
| SRP | **0.806** | 0.806 | 0.743 | 0.693 | 0.488 | 0.700 | 0.682 | 0.218 |
| telomerase | **0.832** | 0.833 | 0.826 | 0.637 | 0.558 | 0.584 | 0.576 | 0.482 |
| RNase P RNA | **0.828** | 0.830 | 0.758 | 0.630 | 0.584 | 0.650 | 0.636 | 0.478 |
| 16S rRNA | 0.620 | 0.623 | 0.584 | 0.622 | **0.663** | 0.511 | 0.469 | 0.605 |
| overall | **0.772** | 0.773 | 0.728 | 0.645 | 0.573 | 0.611 | 0.591 | 0.446 |
| | | | | | F1 scores | | | |
| SRP | **0.804** | 0.804 | 0.779 | 0.695 | 0.486 | 0.681 | 0.677 | 0.323 |
| telomerase | 0.730 | 0.730 | **0.749** | 0.570 | 0.507 | 0.481 | 0.492 | 0.535 |
| RNase P RNA | **0.778** | 0.777 | 0.755 | 0.618 | 0.578 | 0.592 | 0.602 | 0.567 |
| 16S rRNA | 0.617 | 0.617 | 0.596 | 0.603 | **0.662** | 0.488 | 0.466 | 0.616 |
| overall | **0.734** | 0.734 | 0.722 | 0.623 | 0.559 | 0.562 | 0.561 | 0.525 |

| | **Multiple Sequence Alignment Accuracy** | | | | | | | |
|---|---|---|---|---|---|---|---|---|
| | LinearTurboFold (1) | (2) | TurboFold II | LocARNA | MXSCARNA | LinearAlignment | MAFFT | RNAforester (7) |
| | | | | | PPV | | | |
| SRP | **0.463** | 0.463 | 0.458 | 0.305 | 0.387 | 0.414 | 0.393 | 0.263 |
| telomerase | **0.617** | 0.616 | 0.615 | 0.311 | 0.554 | 0.575 | 0.572 | 0.239 |
| RNase P RNA | **0.788** | 0.789 | 0.787 | 0.615 | 0.692 | 0.744 | 0.759 | 0.258 |
| 16S rRNA | 0.971 | 0.977 | **0.977** | 0.647 | 0.971 | 0.947 | 0.974 | 0.239 |
| overall | **0.710** | 0.711 | 0.709 | 0.470 | 0.651 | 0.670 | 0.675 | 0.250 |
| | | | | | Sensitivity | | | |
| SRP | **0.443** | 0.443 | 0.438 | 0.452 | 0.384 | 0.396 | 0.382 | 0.271 |
| telomerase | **0.573** | 0.573 | 0.572 | 0.470 | 0.523 | 0.540 | 0.529 | 0.262 |
| RNase P RNA | **0.765** | 0.766 | **0.765** | 0.596 | 0.684 | 0.724 | 0.738 | 0.286 |
| 16S rRNA | 0.971 | 0.977 | **0.977** | 0.974 | 0.971 | 0.951 | 0.973 | 0.298 |
| overall | **0.688** | 0.690 | **0.688** | 0.623 | 0.641 | 0.653 | 0.656 | 0.280 |
| | | | | | F1 scores | | | |
| SRP | **0.453** | 0.452 | 0.448 | 0.364 | 0.385 | 0.405 | 0.388 | 0.267 |
| telomerase | **0.594** | 0.594 | 0.593 | 0.375 | 0.538 | 0.557 | 0.550 | 0.250 |
| RNase P RNA | **0.776** | 0.777 | **0.776** | 0.605 | 0.688 | 0.734 | 0.748 | 0.271 |
| 16S rRNA | 0.971 | 0.977 | **0.977** | 0.778 | 0.971 | 0.949 | 0.973 | 0.265 |
| overall | **0.699** | 0.700 | 0.698 | 0.535 | 0.646 | 0.661 | 0.665 | 0.264 |

**Table S2. Fully conserved base pairs across 25 complete SARS-CoV-2 and SARS-related genomes with compensatory mutations. The positions and nucleotide type of base pairs correspond to the reference sequence of SARS-CoV-2 (NC_0405512.2). The mutations are from the other 24 genomes.**

| 5' End | 3' End | Base Pair | Average Probability | Mutations | 5' End | 3' End | Base Pair | Average Probability | Mutations |
|---|---|---|---|---|---|---|---|---|---|
| 90 | 121 | GC | 0.971 | AU/GU | 12931 | 12964 | UA | 0.851 | AU/GC |
| 97 | 115 | AU | 0.969 | GU/GC | 13069 | 13108 | UA | 1.000 | CG |
| 153 | 291 | UA | 0.972 | CG | 13078 | 13099 | UA | 1.000 | UG/CG |

Table S2 continued from previous page

| 5' End | 3' End | Base Pair | Average Probability | Mutations | 5' End | 3' End | Base Pair | Average Probability | Mutations |
|---|---|---|---|---|---|---|---|---|---|
| 159 | 282 | GC | 0.961 | AU/GU | 13216 | 13222 | UA | 0.958 | UG/CG |
| 189 | 217 | GC | 0.972 | AU | 13599 | 13628 | UA | 1.000 | CG |
| 358 | 385 | UA | 0.960 | UG/CG | 13638 | 13695 | UA | 0.986 | AU |
| 367 | 373 | CG | 0.961 | UA | 13641 | 13692 | UA | 0.977 | CG |
| 407 | 478 | GC | 0.936 | AU | 14091 | 14107 | UA | 0.989 | CG |
| 442 | 448 | CG | 0.960 | UA/UG | 14161 | 14194 | UA | 1.000 | UG/CG |
| 484 | 555 | UA | 0.877 | AU | 14205 | 14211 | AU | 0.933 | CG |
| 570 | 616 | AU | 0.946 | UA/CG | 14224 | 14251 | AU | 0.996 | GU/GC |
| 652 | 724 | AU | 0.947 | GC | 14355 | 14361 | AU | 0.996 | GC |
| 677 | 703 | GC | 0.933 | AU | 14487 | 14532 | AU | 0.973 | GU/UA/UG/CG |
| 880 | 889 | AU | 0.962 | CG | 14595 | 14604 | UA | 0.999 | UG/CG |
| 970 | 981 | GC | 0.963 | AU | 15435 | 15453 | AU | 0.778 | GC |
| 1231 | 1251 | GC | 0.968 | AU | 15582 | 15607 | AU | 0.993 | GC |
| 1949 | 1956 | UA | 0.929 | UG/CG | 16023 | 16032 | UA | 0.998 | AU |
| 2278 | 2303 | UA | 0.970 | CG | 16080 | 16110 | CG | 0.971 | UA |
| 2855 | 2875 | CG | 0.962 | UA/UG | 16089 | 16101 | GC | 1.000 | AU |
| 2896 | 2923 | UA | 0.973 | AU/GU | 16125 | 16155 | AU | 0.999 | UA |
| 2959 | 2986 | UA | 0.973 | UG/CG | 16230 | 16236 | CG | 0.999 | UA |
| 3712 | 3721 | AU | 0.977 | GC | 16677 | 16716 | GC | 1.000 | AU |
| 3913 | 3928 | UA | 0.979 | AU/UG | 17241 | 17256 | UA | 0.980 | CG |
| 3915 | 3926 | AU | 0.965 | GC | 17244 | 17253 | AU | 0.980 | GC |
| 4096 | 4108 | UA | 0.926 | UG/CG | 17304 | 17331 | CG | 0.981 | UA |
| 4189 | 4225 | CG | 0.980 | GC/UG | 18006 | 18054 | UA | 0.980 | AU/GU |
| 4603 | 4624 | UA | 0.980 | UG/CG | 18439 | 18468 | UA | 0.980 | AU |
| 4978 | 4987 | UA | 0.982 | CG | 18549 | 18561 | AU | 0.982 | CG |
| 5164 | 5203 | GC | 0.975 | AU/GU | 18717 | 18774 | UA | 0.983 | UG/CG |
| 5347 | 5374 | UG | 0.984 | AU/GC/UA | 19074 | 19098 | UA | 0.882 | AU |
| 5356 | 5371 | UA | 0.953 | AU/GC | 19386 | 19419 | CG | 0.986 | UA/UG |
| 5417 | 5428 | UA | 0.982 | AU | 19395 | 19410 | UA | 0.981 | CG |
| 5476 | 5521 | AU | 0.940 | GU/GC | 19707 | 19732 | CG | 0.981 | UA |
| 5482 | 5515 | CG | 0.984 | UA/UG | 19708 | 19731 | AU | 0.985 | GU/GC |
| 5739 | 5770 | GC | 0.984 | AU | 19917 | 19953 | UA | 0.980 | AU |
| 6034 | 6055 | AU | 0.983 | GC | 19929 | 19941 | UA | 0.984 | AU |
| 6037 | 6052 | CG | 0.982 | UA | 20172 | 20187 | UA | 0.977 | CG |
| 6154 | 6202 | AU | 0.987 | GC | 20217 | 20265 | UA | 0.939 | CG |
| 6328 | 6343 | AU | 0.988 | UA | 20223 | 20260 | AU | 0.981 | GC |
| 6364 | 6388 | GC | 0.989 | AU | 20523 | 20541 | UA | 0.988 | CG |
| 6367 | 6385 | GC | 0.989 | AU | 20841 | 20901 | AU | 0.988 | GU/GC |
| 6458 | 6490 | AU | 0.988 | GC | 20985 | 20997 | AU | 0.757 | UA |
| 6460 | 6488 | UA | 0.988 | CG | 21163 | 21201 | AU | 0.980 | GU/GC |
| 6903 | 6922 | CG | 0.895 | UA | 21300 | 21321 | AU | 0.988 | GU/GC |
| 6977 | 7006 | GC | 0.970 | AU/GU | 21411 | 21423 | CG | 0.989 | UA |
| 7103 | 7135 | AU | 0.891 | GU/GC | 21513 | 21523 | CG | 0.988 | UA |
| 7480 | 7531 | UA | 0.942 | UG/CG | 22837 | 22903 | AU | 0.988 | GU/GC |
| 7558 | 7597 | AU | 0.956 | GC | 23531 | 23548 | AU | 0.717 | GC |
| 7864 | 7876 | AU | 0.972 | GU/GC | 23621 | 23647 | GC | 0.991 | AU |
| 8146 | 8219 | CG | 0.993 | UA | 23797 | 23806 | UA | 0.931 | UG/CG |
| 8147 | 8218 | AU | 0.992 | GU/GC | 23980 | 24088 | AU | 0.978 | GU/GC |
| 8153 | 8212 | UA | 0.987 | CG | 23983 | 24085 | AU | 0.974 | UA/CG |
| 8317 | 8332 | AU | 0.995 | GU/GC | 24121 | 24152 | AU | 0.994 | GC |
| 8437 | 8458 | AU | 0.915 | GU/GC | 24553 | 24586 | CG | 0.996 | UA |
| 8698 | 8738 | UA | 0.824 | CG | 24757 | 24766 | GC | 0.974 | GU/CG |
| 8860 | 8881 | CG | 0.996 | AU/UA | 25336 | 25370 | AU | 0.906 | GU/GC |

**Table S2 continued from previous page**

| 5' End | 3' End | Base Pair | Average Probability | Mutations | 5' End | 3' End | Base Pair | Average Probability | Mutations |
|---|---|---|---|---|---|---|---|---|---|
| 9046 | 9079 | UA | 0.995 | UG/CG | 25991 | 26004 | GC | 0.996 | AU/GU |
| 9055 | 9070 | AU | 0.969 | GC | 26145 | 26190 | UA | 0.997 | CG |
| 9427 | 9433 | UA | 0.991 | CG | 26262 | 26305 | GC | 0.911 | AU |
| 9472 | 9511 | AU | 0.996 | UA | 26630 | 26658 | AU | 0.903 | GC |
| 9689 | 9703 | AU | 0.932 | GC | 26676 | 26706 | AU | 0.986 | GC |
| 9842 | 9874 | UA | 0.990 | UG/CG | 26939 | 26975 | AU | 0.996 | CG |
| 10213 | 10248 | UA | 0.997 | CG | 27412 | 27456 | UA | 0.998 | UG/CG |
| 10651 | 10669 | UA | 0.998 | CG | 27415 | 27453 | GC | 0.994 | AU |
| 10864 | 10906 | AU | 0.926 | GC | 27603 | 27613 | CG | 0.996 | UA |
| 10873 | 10898 | AU | 0.946 | GC | 27699 | 27744 | UA | 0.935 | CG |
| 10984 | 11026 | AU | 0.976 | GC | 27717 | 27725 | GC | 0.990 | AU/GU |
| 11782 | 11803 | AU | 0.981 | GU/GC | 28642 | 28664 | UA | 0.997 | UG/CG |
| 11788 | 11797 | CG | 1.000 | UA | 28910 | 28930 | AU | 0.997 | GU/GC |
| 11971 | 12013 | AU | 0.966 | GU/GC | 29567 | 29597 | AU | 0.999 | GC |
| 11989 | 11995 | UA | 0.763 | UG/CG | 29635 | 29651 | CG | 0.999 | AU |
| 12538 | 12577 | UA | 0.940 | UG/CG | 29637 | 29649 | UA | 0.991 | CG |

**Table S3. Fully conserved structures among 25 genomes. Regions with compensatory mutations are annotated with alternative base pairs. Novel regions compared with Rangan *et al.* (20, 21) are annotated with stars.**

| Region | Sequence & Structure | Compensatory Mutations |
|---|---|---|
| 45-59 | GAUCUCUUGUAGAUC<br>(((((.....))))) | |
| 84-127 | CUGUGUGGCUGUCACUCGGCUGCAUGCUUAGUGCACUCACGCAG<br>((((((((.((.((((.(((.....))).)))))).)))))))) | GC ->AU<br>AU ->GC |
| 626-643* | GUUCUUCUUCGUAAGAAC<br>((((((......)))))) | |
| 1324-1341 | UGCCACUACUUGUGGUUA<br>((((((.....))))).) | |
| 1685-1698 | GCCAUUAUUUUGGC<br>((((......)))) | |
| 1785-1800* | GUAAUUUUAAAGUUAC<br>((((((....)))))) | |
| 2818-2837* | AGUACUUAAUGAGAAGUGCU<br>(((((((......))))))) | |
| 4973-4993 | GUGUUUACAACAGUAGACAAC<br>((((((((....))))))).) | UA ->CG |
| 5021-5033 | AUGUCAAUGACAU<br>(((((...))))) | |
| 6450-6498 | UUGAGUGUAAUGUGAAAACUACCGAAGUUGUAGGAGACAUUAUACUUAA<br>(((((((((((((.....((((.......))))...))))))))))))) | AU ->GC<br>UA ->CG |
| 8078-8084 | CCAAUGG<br>((...)) | |
| 10494-10509* | GUGUUGGUUUUAACAU<br>((((((....)))))) | |
| 11131-11140 | UGCUUUUGCA<br>(((....))) | |
| 12203-12218* | UUGAAGAAGUCUUUGA<br>((((((....)))))) | |
| 12257-12268 | CAACGUAAGUUG<br>((((....)))) | |
| 12386-12412 | GAUAAUGAUGCACUCAACAACAUUAUC<br>(((((((.((.......)).))))))) | |
| 12672-12685 | CUGUCAAAUUACAG<br>((((......)))) | |
| 12904-12926* | UAGGUUUGUUACAGACACACCUA<br>(((((.((((...)))).))))) | |
| 12970-12988* | CAACCUAAAUAGAGGUAUG<br>(((((......)))).)) | |
| 13409-13422* | AGUUGUGAUCAACU<br>(((((....))))) | |
| 14729-14769 | AGGAAGGAAGUUCUGUUGAAUUAAAACACUUCUUCUUUGCU<br>((.(((((((...((((.......))))...))))))).)) | |
| 14773-14790 | GAUGGUAAUGCUGCUAUC<br>(((((((....))))))) | |
| 14794-14818 | GAUUAUGACUACUAUCGUUAUAAUC<br>(((((((((.......))))))))) | |
| 15254-15269* | UUUAUAGUGAUGUAGA<br>((((((....)))))) | |
| 15430-15458* | AGUGAAAUGGUCAUGUGUGGCGGUUCACU<br>((((((.(.((((....)))).))))))) | AU ->GC |
| 15502-15509* | GCUUAUGC<br>((....)) | |
| 15618-15628 | ACUUUAUGAGU<br>((((...)))) | |
| 15775-15801* | AUAAAGAACUUUAAGUCAGUUCUUUAU<br>((((((((((.......)))))))))) | |
| 16013-16042* | GGUUCGUGUCUUUAGCUAUAGAUGCUUACC<br>(((..((((((........))))))..))) | UA ->AU |
| 16180-16194 | AGGUAUUGGGAACCU<br>((((.......)))) | |
| 16955-16973* | UAGUGCCACAAGAGCACUA<br>(((((((....).)))))) | |

**Table S3 continued from previous page**

| Region | Sequence & Structure | Compensatory Mutations |
|---|---|---|
| 17236-17262 | AUACCUGCACGUGCUCGUGUAGAGUGU<br>(((((((((((....))))))).)))) | UA ->CG<br>AU ->GC |
| 17462-17470 | GCACAUUGC<br>(((...))) | |
| 18547-18563* | UUAAGUGACACACUUAA<br>(((((((...))))))) | AU ->CG |
| 18660-18674* | UUGUCUAUGUGAUAG<br>(((((.....))))) | |
| 18848-18872* | CUAGUUGUGAUGCAAUCAUGACUAG<br>(((((((((((...))))))))))) | |
| 19036-19043* | GGUAACCC<br>((....)) | |
| 19973-20002* | CACUCACUGUCUUUUUUGAUGGUAGAGUUG<br>(((((((((((......)))))).)))).) | |
| 20103-20127* | UCUUAAUGGAGUCACAUUAAUUGGA<br>((((((((......)))))...))) | |
| 20709-20719* | GCAAAGAAUGC<br>(((.....))) | |
| 23101-23110* | UUCUUUUGAA<br>(((....))) | |
| 23119-23128 | UGCACCAGCA<br>(((....))) | |
| 23796-23807 | AUUCAACUGAAU<br>((((....)))) | UA ->CG |
| 24305-24319 | GUUCUCUAUGAGAAC<br>((((((...)))))) | |
| 25380-25407* | CAUAAACGAACUUAUGGAUUUGUUUAUG<br>((((((((((((...)).)))))))))) | |
| 25937-25959 | AUGACUACCAGAUUGGUGGUUAU<br>((((((((((...)))))))))) | |
| 26209-26224 | GUGCCUUUGUAAGCAC<br>((((........)))) | |
| 26358-26379 | AUUGUGUGCGUACUGCUGCAAU<br>((((((.(((...))))))))) | |
| 26581-26600 | GGAACCUAGUAAUAGGUUUC<br>((((((((....)))))))) | |
| 26713-26740 | GUUUUGUGCUUGCUGCUGUUUACAGAAU<br>((((((((...((....)).)))))))) | |
| 27635-27651* | CAGUUUCACCUAAACUG<br>((((((.....)))))) | |
| 28755-28766 | UUCCUCAAGGAA<br>((((....)))) | |
| 29145-29166 | UAAUCAGACAAGGAACUGAUUA<br>(((((((........))))))) | |
| 29210-29225 | GCGUUCUUCGGAAUGU<br>((((((....)))))) | |
| 29240-29253 | GAAGUCACACCUUC<br>((((......)))) | |
| 29288-29306 | UUGGAUGACAAAGAUCCAA<br>((((((.......)))))) | |
| 29321-29347 | GUCAUUUUGCUGAAUAAGCAUAUUGAC<br>((((...((((.....))))...)))) | |
| 29470-29488* | UUUGGAUGAUUUCUCCAAA<br>((((((.(....))))))) | |
| 29548-29613 | CACAAGGCAGAUGGGCUAUAUAAACGUUUUCGCUUUUCCGUUUACGAUAUAUAGUCUACUCUUGUG<br>((((((..((.(((((((((((..(((...((......))...))).))))))))))))))))))) | UA ->CG |

**Table S4. Accessibility and conservation of target regions for public RT-PCR forward/reverse primers and probes (22). The accessibility is computed by LinearTurboFold, and it is underlined if larger than zero. The conservation on 9 SARS-related genomes is the number of mutated sites. The conservation on the $\sim$2M SARS-CoV-2 dataset is the percentage of exact matches, which is underlined or bold if less than 0.97 or 0.5, respectively. (The average sequence identity of SARS-CoV-2 genomes is 0.9987, and the average length of primers and probes is 23 $nt$. Therefore, the probability of randomly sampling a region of length 23 $nt$ without mutations is $0.9987^{23} \approx 0.97$).**

| Gene | Institute | Forward Primer / Probe / Reverse Primer | | | | |
|---|---|---|---|---|---|---|
| | | Start | Length | Accessibility | Conservation: SARS-related # mut. site | Conservation: SARS-CoV-2 (2M) exact match |
| ORF1ab nsp9 | Institut Pasteur (1) | 12689 / 12717 / 12779 | 18 / 21 / 18 | 0.0000 / 0.0000 / <u>0.0160</u> | 4 / 3 / 5 | 0.9989 / 0.9967 / 0.9829 |
| ORF1ab nsp10 | China CDC (1) | 13341 / 13377 / 13441 | 21 / 30 / 19 | 0.0000 / 0.0000 / 0.0000 | 4 / 3 / 11 | 0.9937 / 0.9875 / 0.9868 |
| ORF1ab nsp12 (RdRp) | Institut Pasteur (2) | 14079 / 14105 / 14166 | 19 / 19 / 20 | 0.0000 / <u>0.0001</u> / 0.0000 | 4 / 8 / 9 | 0.9978 / 0.9332 / 0.9941 |
| | Charite Germany (1) | 15430 / 15469 / 15504 | 22 / 25 / 26 | 0.0000 / 0.0000 / 0.0000 | 0 / 6 / 1 | <u>0.9167</u> / 0.9938 / 0.9982 |
| ORF1ab nsp14 | HKU (1) | 18777 / 18849 / 18888 | 20 / 24 / 21 | 0.0000 / 0.0000 / 0.0000 | 1 / 1 / 3 | 0.9958 / 0.9969 / 0.9933 |
| E | Charite Germany (2) | 26268 / 26332 / 26359 | 26 / 26 / 22 | 0.0000 / 0.0000 / 0.0000 | 0 / 0 / 0 | 0.9958 / 0.9969 / 0.9933 |
| N | CDC (1) | 28286 / 28309 / 28334 | 20 / 24 / 24 | 0.0000 / 0.0000 / 0.0000 | 12 / 2 / 8 | 0.9913 / 0.9762 / 0.9934 |
| | NIH Thailand | 28319 / 28341 / 28357 | 20 / 16 / 19 | 0.0000 / <u>0.0026</u> / 0.0000 | 3 / 6 / 6 | 0.9908 / 0.9953 / 0.9927 |
| | CDC (2) | 28680 / 28704 / 28731 | 22 / 24 / 21 | 0.0000 / 0.0000 / <u>0.0010</u> | 4 / 4 / 2 | 0.9862 / 0.9796 / 0.9895 |
| | Charite Germany (3) | 28705 / 28753 / 28813 | 19 / 25 / 20 | 0.0000 / <u>0.0003</u> / 0.0000 | 2 / 0 / 6 | 0.9914 / 0.9920 / 0.9858 |
| | China CDC (2) | 28880 / 28934 / 28957 | 22 / 20 / 22 | <u>0.0710</u> / 0.0000 / 0.0000 | 5 / 7 / 4 | **0.2734** / 0.9911 / **0.4844** |
| | NIID Japan | 29124 / 29222 / 29262 | 20 / 20 / 20 | 0.0000 / 0.0000 / 0.0000 | 6 / 1 / 7 | 0.9953 / 0.9785 / 0.9853 |
| | HKU (2) | 29144 / 29179 / 29235 | 22 / 20 / 19 | 0.0000 / 0.0000 / 0.0000 | 2 / 2 / 1 | 0.9904 / 0.9945 / 0.9895 |
| | CDC (3) | 29163 / 29188 / 29212 | 20 / 23 / 18 | 0.0000 / 0.0000 / 0.0000 | 2 / 6 / 2 | 0.9892 / 0.9797 / 0.9901 |

**Table S5. Accessible and conserved regions with two kinds of constraints on conservation: 1) at most three mutations on SARS-related genomes; 2) the average sequence identity on the SARS-CoV-2 dataset at least 0.999. The start positions and sequences correspond to the reference sequence of SARS-CoV-2 (NC_0405512.2). The accessibilities are calculated from folding with homologs (LinearTurboFold) and single sequence folding (LinearPartition), respectively. We searched for these regions among human representative transcript set (RefSeq Select RNA sequences, refseq_select) using BLAST, and several regions have the exact matches with human transcripts (underlined). Using single sequence folding can only get one accessible region (bold). The conservation of these regions on 9 SARS-related genomes is the number of mutated sites. The table also shows two types of conservations on a large SARS-CoV-2 dataset containing $\sim$2M genomes submitted to GISAID up to June 30, 2021: the average sequence identity with reference sequence, and the percentage of exact matches of the whole region, respectively.**

| Region | Start | Length | Sequence | Gene | Accessibility | | | | Conservation | | BLAST Match | GC (%) |
|---|---|---|---|---|---|---|---|---|---|---|---|---|
| | | | | | LinearTurboFold (Homologous Folding) | | | Single Seq. | SARSr (9) | SARS-CoV-2 (2M) | | |
| | | | | | Average | Range | $\Delta G$(kcal/mol) | Folding Range | # Mut. Sites | Identity / Exact | | |
| Region 1 | 739 | 18 | AGAAAACUGGAACACUAA | ORF1ab nsp1 | 0.71 ± 0.04 | 0.62 – 0.76 | 0.22 ± 0.04 | 0.00 – 0.00 | 2/18 | 0.9998 / 0.9970 | 14/18 | 33 |
| Region 2 | 995 | 17 | CGUUCUGAAAAGAGCUA | | 0.99 ± 0.00 | 0.99 – 0.99 | 0.01 ± 0.00 | 0.00 – 0.00 | 3/17 | 0.9999 / 0.9985 | 14/17 | 41 |
| Region 3 | 998 | 17 | UCUGAAAAGAGCUAUGA | ORF1ab nsp2 | 1.00 ± 0.00 | 1.00 – 1.00 | 0.00 ± 0.00 | 0.00 – 0.00 | 3/17 | 0.9999 / 0.9984 | 15/17 | 35 |
| Region 4 | 1001 | 15 | GAAAAGAGCUAUGAA | | 0.74 ± 0.08 | 0.51 – 0.80 | 0.19 ± 0.07 | 0.00 – 0.00 | 3/15 | 0.9999 / 0.9985 | 15/15 | 33 |
| Region 5 | 6765 | 16 | AUUAUAUGCCUUAUUU | | 0.96 ± 0.01 | 0.95 – 0.97 | 0.02 ± 0.00 | 0.30 – 0.40 | 3/16 | 1.0000 / 0.9993 | 14/16 | 19 |
| Region 6 | 6767 | 15 | UAUAUGCCUUAUUUC | ORF1ab nsp3 (PLpro) | 0.96 ± 0.01 | 0.95 – 0.97 | 0.02 ± 0.00 | 0.37 – 0.42 | 3/15 | 0.9999 / 0.9981 | 15/15 | 27 |
| Region 7 | 7691 | 22 | CAGUUUAAAAGACCAAUAAAUC | | 0.77 ± 0.03 | 0.69 – 0.83 | 0.16 ± 0.03 | 0.00 – 0.00 | 3/22 | 1.0000 / 0.9991 | 20/22 | 27 |
| Region 8 | 9527 | 18 | UCAUUCACUGUACUCUGU | | 0.66 ± 0.03 | 0.60 – 0.70 | 0.26 ± 0.03 | 0.00 – 0.00 | 3/18 | 0.9997 / 0.9945 | 15/18 | 39 |
| Region 9 | 9530 | 17 | UUCACUGUACUCUGUUU | ORF1ab nsp4 | 0.64 ± 0.03 | 0.57 – 0.68 | 0.28 ± 0.03 | 0.00 – 0.00 | 3/17 | 0.9998 / 0.9965 | 15/17 | 35 |
| Region 10 | 9905 | 15 | UACAAGUAUUUUAGU | | 0.75 ± 0.10 | 0.51 – 0.82 | 0.18 ± 0.10 | 0.00 – 0.00 | 2/15 | 0.9999 / 0.9980 | 15/15 | 20 |
| Region 11 | 10010 | 17 | CUUUACCAACCACCACA | | 0.75 ± 0.06 | 0.54 – 0.79 | 0.18 ± 0.06 | 0.00 – 0.01 | 3/17 | 0.9999 / 0.9989 | 15/17 | 47 |
| Region 12 | 11536 | 25 | UAUUGUUUUUAUGUGUGUUGAGUAU | | 0.67 ± 0.06 | 0.59 – 0.78 | 0.25 ± 0.05 | 0.00 – 0.01 | 3/25 | 0.9998 / 0.9961 | 17/25 | 24 |
| Region 13 | 11540 | 22 | GUUUUUAUGUGUGUUGAGUAUU | ORF1ab nsp6 | 0.79 ± 0.08 | 0.69 – 0.92 | 0.15 ± 0.06 | 0.00 – 0.02 | 3/22 | 0.9998 / 0.9965 | 17/22 | 27 |
| Region 14 | 11543 | 20 | UUUAUGUGUGUUGAGUAUUG | | 0.78 ± 0.08 | 0.69 – 0.92 | 0.15 ± 0.06 | 0.00 – 0.00 | 3/20 | 0.9998 / 0.9970 | 17/20 | 30 |
| Region 15 | 11547 | 19 | UGUGUGUUGAGUAUUGCCC | | 0.79 ± 0.08 | 0.69 – 0.92 | 0.15 ± 0.06 | 0.00 – 0.00 | 3/19 | 0.9998 / 0.9954 | 15/19 | 47 |
| Region 16 | 13454 | 15 | CAAUCGUUUUUAAAC | ORF1ab nsp11 | 0.96 ± 0.04 | 0.88 – 0.98 | 0.02 ± 0.02 | 0.07 – 0.11 | 3/15 | 0.9998 / 0.9972 | 14/15 | 27 |
| Region 17 | 15141 | 22 | CAAUAGACAGUUUCAUCAAAAA | | 0.61 ± 0.05 | 0.50 – 0.67 | 0.30 ± 0.06 | 0.00 – 0.00 | 3/22 | 0.9999 / 0.9986 | 18/22 | 27 |
| Region 18 | 15890 | 15 | CAAUGCUAGUUAAAC | ORF1ab nsp12 (RdRp) | 0.63 ± 0.06 | 0.50 – 0.68 | 0.29 ± 0.06 | 0.00 – 0.23 | 1/15 | 0.9999 / 0.9991 | 13/15 | 33 |
| Region 19 | 15997 | 16 | ACACUUAUGAUUGAAC | | 0.72 ± 0.03 | 0.65 – 0.76 | 0.20 ± 0.03 | 0.00 – 0.40 | 2/16 | 1.0000 / 0.9997 | 13/16 | 31 |
| Region 20 | 17194 | 22 | AAGGCAUUAAAAUAUUUGCCUA | | 1.00 ± 0.00 | 0.99 – 1.00 | 0.00 ± 0.00 | 0.00 – 0.00 | 3/22 | 0.9999 / 0.9989 | 16/22 | 27 |
| Region 21 | 18032 | 17 | CUUUACAAGCUGAAAAU | ORF1ab nsp13 (helicase) | 0.67 ± 0.05 | 0.57 – 0.73 | 0.25 ± 0.05 | 0.00 – 0.00 | 2/17 | 0.9999 / 0.9978 | 14/17 | 29 |
| Region 22 | 18035 | 15 | UACAAGCUGAAAAUG | | 0.91 ± 0.10 | 0.54 – 0.95 | 0.07 ± 0.08 | 0.00 – 0.02 | 1/15 | 1.0000 / 0.9993 | 13/15 | 33 |
| Region 23 | 18036 | 17 | ACAAGCUGAAAAUGUAA | | 0.93 ± 0.09 | 0.57 – 0.97 | 0.05 ± 0.08 | 0.00 – 0.02 | 2/17 | 0.9998 / 0.9992 | 15/17 | 29 |
| Region 24 | 20134 | 20 | GUAAAAACACAGUUCAAUUA | ORF1ab nsp15 | 0.62 ± 0.05 | 0.52 – 0.68 | 0.29 ± 0.05 | 0.00 – 0.07 | 3/20 | 0.9998 / 0.9959 | 14/20 | 25 |
| Region 25 | 20135 | 21 | UAAAAACACAGUUCAAUUAUU | | 0.63 ± 0.04 | 0.54 – 0.68 | 0.29 ± 0.04 | 0.00 – 0.07 | 3/21 | 0.9998 / 0.9967 | 14/21 | 19 |
| Region 26 | 25546 | 17 | CUUCUUGCUGUUUUUCA | ORF3a | 0.94 ± 0.02 | 0.89 – 0.95 | 0.04 ± 0.01 | 0.00 – 0.02 | 1/17 | 0.9998 / 0.9904 | 16/17 | 35 |
| **Region 27** | **27132** | **15** | **UAUAAAUUAAACACA** | **M** | **0.70 ± 0.01** | **0.68 – 0.72** | **0.22 ± 0.01** | **0.65 – 0.79** | 3/15 | 0.9998 / 0.9963 | 15/15 | 13 |
| Region 28 | 27525 | 16 | ACCAUUUCAUCCUCUA | ORF7a | 0.99 ± 0.00 | 0.99 – 1.00 | 0.00 ± 0.00 | 0.00 – 0.05 | 3/16 | 0.9993 / 0.9927 | 14/16 | 38 |
| Region 29 | 28402 | 16 | AGGUUUACCCAAUAAU | | 1.00 ± 0.00 | 0.99 – 1.00 | 0.00 ± 0.00 | 0.00 – 0.81 | 1/16 | 0.9999 / 0.9985 | 14/16 | 31 |
| Region 30 | 28690 | 15 | GAAUACACCAAAAGA | | 0.78 ± 0.01 | 0.77 – 0.78 | 0.16 ± 0.00 | 0.01 – 0.52 | 3/33 | 0.9992 / 0.9879 | 14/15 | 33 |
| Region 31 | 28691 | 20 | AAUACACCAAAAGAUCACAU | N | 0.77 ± 0.01 | 0.76 – 0.78 | 0.16 ± 0.00 | 0.00 – 0.52 | 3/20 | 0.9994 / 0.9883 | 15/20 | 30 |
| Region 32 | 28694 | 18 | ACACCAAAAGAUCACAUU | | 0.77 ± 0.01 | 0.76 – 0.78 | 0.16 ± 0.00 | 0.01 – 0.52 | 3/18 | 0.9994 / 0.9886 | 16/18 | 33 |
| Region 33 | 29075 | 15 | UACAAUGUAACACAA | | 1.00 ± 0.00 | 1.00 – 1.00 | 0.00 ± 0.00 | 0.01 – 0.83 | 3/15 | 0.9996 / 0.9942 | 12/15 | 27 |

**Table S6. Accessible and conserved regions with a loose constraint on conservation: the average sequence identity on the ∼2M SARS-CoV-2 dataset is at least 0.999. The table keeps the same format as Tab. S5 and only displays new regions not included in that table.**

| Region | Start | Length | Sequence | Gene | Accessibility | | | | Conservation | | BLAST | GC |
|---|---|---|---|---|---|---|---|---|---|---|---|---|
| | | | | | LinearTurboFold (Homologous Folding) | | | Single Seq. | SARSr (9) | SARS-CoV-2 (2M) | Match | (%) |
| | | | | | Average | Range | $\Delta G$(kcal/mol) | Folding Range | # Mut. Sites | Identity / Exact | | |
| Region 1 | 1094 | 19 | UUAAAUUCCAUAAUCAAGA | | 0.86 $\pm$ 0.03 | 0.80 – 0.89 | 0.10 $\pm$ 0.02 | 0.01 – 0.10 | 8/19 | 0.9998 / 0.9963 | 17/19 | 21 |
| Region 2 | 1301 | 19 | ACUGAGAAUUUGACUAAAG | | 0.75 $\pm$ 0.05 | 0.64 – 0.79 | 0.18 $\pm$ 0.04 | 0.00 – 0.00 | 8/19 | 0.9997 / 0.9950 | 14/19 | 32 |
| Region 3 | 1359 | 18 | UUGUUAAAAUUUAUUGUC | ORF1ab nsp2 | 0.75 $\pm$ 0.02 | 0.72 – 0.81 | 0.17 $\pm$ 0.02 | 0.09 – 0.22 | 7/18 | 0.9999 / 0.9989 | 16/18 | 17 |
| Region 4 | 1420 | 18 | CGAAUACCAUAAUGAAUC | | 0.94 $\pm$ 0.00 | 0.94 – 0.95 | 0.04 $\pm$ 0.00 | 0.01 – 0.03 | 7/18 | 0.9997 / 0.9941 | 13/18 | 33 |
| Region 5 | 2550 | 19 | AUUUACAACCAUUAGAACA | | 0.93 $\pm$ 0.03 | 0.89 – 0.96 | 0.04 $\pm$ 0.02 | 0.24 – 0.31 | 12/19 | 0.9998 / 0.9971 | 13/19 | 26 |
| Region 6 | 3648 | 15 | UUCAACUUCUUAAGA | | 0.93 $\pm$ 0.02 | 0.91 – 0.96 | 0.04 $\pm$ 0.01 | 0.00 – 0.03 | 8/15 | 0.9999 / 0.9980 | 14/15 | 27 |
| Region 7 | 3733 | 19 | UGACCCUAUACAUUCUUUA | | 0.91 $\pm$ 0.01 | 0.89 – 0.91 | 0.06 $\pm$ 0.00 | 0.00 – 0.01 | 8/19 | 0.9996 / 0.9928 | 13/19 | 32 |
| Region 8 | 4405 | 17 | ACAUGCAGAAGAAACAC | ORF1ab nsp3 | 0.55 $\pm$ 0.02 | 0.51 – 0.59 | 0.36 $\pm$ 0.02 | 0.00 – 0.00 | 6/17 | 0.9999 / 0.9987 | 15/17 | 41 |
| Region 9 | 4406 | 21 | CAUGCAGAAGAAACACGCAAA | (PLpro) | 0.75 $\pm$ 0.03 | 0.71 – 0.80 | 0.18 $\pm$ 0.02 | 0.00 – 0.00 | 7/21 | 0.9999 / 0.9975 | 17/21 | 43 |
| Region 10 | 4864 | 26 | AAGUGUAUAUUACACUAGUAAUCCUA | | 0.85 $\pm$ 0.07 | 0.60 – 0.88 | 0.11 $\pm$ 0.06 | 0.00 – 0.00 | 14/26 | 0.9999 / 0.9975 | 16/26 | 27 |
| Region 11 | 5773 | 23 | UAAACAUAUAACUUCUAAAGAAA | | 0.64 $\pm$ 0.04 | 0.57 – 0.70 | 0.28 $\pm$ 0.04 | 0.00 – 0.21 | 9/23 | 0.9998 / 0.9961 | 16/23 | 17 |
| Region 12 | 6129 | 16 | UUAAAGUUACAUUUUU | | 0.97 $\pm$ 0.04 | 0.79 – 0.98 | 0.02 $\pm$ 0.03 | 0.01 – 0.11 | 6/16 | 1.0000 / 0.9996 | 16/16 | 13 |
| Region 13 | 6499 | 32 | ACCAGCAAAUAAUAGUUUAAAAAUUACAGAAG | | 0.61 $\pm$ 0.02 | 0.55 – 0.63 | 0.31 $\pm$ 0.03 | 0.00 – 0.00 | 15/32 | 0.9997 / 0.9941 | 19/32 | 25 |
| Region 14 | 6622 | 19 | GAAAACCCUUGCUACUCAU | | 0.95 $\pm$ 0.00 | 0.94 – 0.96 | 0.03 $\pm$ 0.00 | 0.00 – 0.01 | 8/19 | 0.9993 / 0.9869 | 15/19 | 42 |
| Region 15 | 6697 | 15 | UUUUCUUAACAAAGU | | 0.96 $\pm$ 0.00 | 0.95 – 0.97 | 0.03 $\pm$ 0.00 | 0.00 – 0.00 | 11/15 | 0.9995 / 0.9920 | 14/15 | 20 |
| Region 16 | 7010 | 15 | GCUUUAGGUGUUUUA | | 0.76 $\pm$ 0.03 | 0.69 – 0.79 | 0.17 $\pm$ 0.02 | 0.00 – 0.01 | 7/15 | 0.9999 / 0.9982 | 14/15 | 33 |
| Region 17 | 7073 | 20 | UAUUUGAACUCUACUAAUGU | | 0.85 $\pm$ 0.02 | 0.83 – 0.88 | 0.10 $\pm$ 0.01 | 0.00 – 0.24 | 7/20 | 0.9998 / 0.9967 | 15/20 | 25 |
| Region 18 | 7725 | 19 | CUUCUUACAUCGUUGAUAG | | 0.76 $\pm$ 0.06 | 0.69 – 0.86 | 0.17 $\pm$ 0.05 | 0.00 – 0.00 | 8/19 | 0.9996 / 0.9930 | 15/19 | 37 |
| Region 19 | 9336 | 15 | UAAAUUUACUUACUA | ORF1ab nsp4 | 1.00 $\pm$ 0.00 | 1.00 – 1.00 | 0.00 $\pm$ 0.00 | 0.00 – 0.95 | 8/15 | 0.9998 / 0.9976 | 13/15 | 13 |
| Region 20 | 9555 | 15 | UUUACUCAUUCUUAC | | 0.89 $\pm$ 0.01 | 0.88 – 0.91 | 0.07 $\pm$ 0.01 | 0.00 – 0.62 | 9/15 | 0.9995 / 0.9927 | 13/15 | 27 |
| Region 21 | 11629 | 18 | UUUUUGUACUUGUUACUU | ORF1ab nsp6 | 0.77 $\pm$ 0.07 | 0.69 – 0.88 | 0.16 $\pm$ 0.05 | 0.00 – 0.27 | 8/18 | 0.9998 / 0.9981 | 14/18 | 22 |
| Region 22 | 12825 | 17 | AUUUACAGGAUUUGAAA | | 0.75 $\pm$ 0.06 | 0.62 – 0.82 | 0.18 $\pm$ 0.05 | 0.00 – 0.01 | 9/17 | 0.9999 / 0.9991 | 16/17 | 24 |
| Region 23 | 14170 | 16 | AUAUUAACCUUGACCA | ORF1ab nsp12 (RdRp) | 0.90 $\pm$ 0.01 | 0.88 – 0.92 | 0.06 $\pm$ 0.01 | 0.00 – 0.27 | 7/16 | 0.9997 / 0.9948 | 14/16 | 31 |
| Region 24 | 16339 | 18 | AUAUCAACAUCACAUAAA | ORF1ab nsp13 | 0.96 $\pm$ 0.06 | 0.83 – 1.00 | 0.02 $\pm$ 0.04 | 0.00 – 0.13 | 5/18 | 0.9999 / 0.9988 | 15/18 | 22 |
| Region 25 | 17651 | 16 | UUAAAAUGUUUUAUAA | (helicase) | 0.96 $\pm$ 0.00 | 0.95 – 0.97 | 0.02 $\pm$ 0.00 | 0.00 – 0.17 | 5/16 | 0.9999 / 0.9988 | 15/16 | 6 |
| Region 26 | 20652 | 16 | AUUACAAUCUAGUCAA | ORF1ab nsp15 | 0.73 $\pm$ 0.02 | 0.70 – 0.77 | 0.19 $\pm$ 0.02 | 0.00 – 0.38 | 6/16 | 0.9995 / 0.9918 | 13/16 | 25 |
| Region 27 | 20844 | 24 | CUAUAAUAUGAGAGUUAUACAUUU | ORF1ab nsp16 | 1.00 $\pm$ 0.00 | 1.00 – 1.00 | 0.00 $\pm$ 0.00 | 0.00 – 0.02 | 7/24 | 0.9999 / 0.9982 | 14/24 | 21 |
| Region 28 | 21622 | 16 | CAGAACUCAAUUACCC | | 0.81 $\pm$ 0.03 | 0.78 – 0.89 | 0.13 $\pm$ 0.02 | 0.00 – 0.36 | 15/16 | 0.9992 / 0.9872 | 13/16 | 44 |
| Region 29 | 21922 | 16 | UAAUAACGCUACUAAU | | 0.96 $\pm$ 0.01 | 0.93 – 0.97 | 0.03 $\pm$ 0.01 | 0.00 – 0.00 | 7/16 | 0.9999 / 0.9988 | 14/16 | 25 |
| Region 30 | 21950 | 15 | GUCUGUGAAUUUCAA | | 0.73 $\pm$ 0.05 | 0.67 – 0.82 | 0.19 $\pm$ 0.04 | 0.00 – 0.11 | 6/15 | 0.9999 / 0.9983 | 14/15 | 33 |
| Region 31 | 22876 | 24 | UAACAAUCUUGAUUCUAAGGUUGG | | 0.94 $\pm$ 0.04 | 0.86 – 0.97 | 0.04 $\pm$ 0.02 | 0.00 – 0.00 | 23/24 | 0.9993 / 0.9825 | 16/24 | 33 |
| Region 32 | 23031 | 16 | UUCCUUUACAAUCAUA | S | 0.98 $\pm$ 0.00 | 0.98 – 0.98 | 0.01 $\pm$ 0.00 | 0.00 – 0.04 | 14/16 | 0.9995 / 0.9920 | 13/16 | 25 |
| Region 33 | 24058 | 15 | CUUCAUCAAACAAUA | | 0.75 $\pm$ 0.13 | 0.51 – 0.87 | 0.19 $\pm$ 0.12 | 0.00 – 0.00 | 5/15 | 0.9999 / 0.9988 | 14/15 | 27 |
| Region 34 | 24166 | 19 | AAUGAUUGCUCAAUACACU | | 0.73 $\pm$ 0.06 | 0.61 – 0.78 | 0.20 $\pm$ 0.05 | 0.00 – 0.16 | 8/19 | 0.9999 / 0.9977 | 17/19 | 32 |
| Region 35 | 24170 | 16 | AUUGCUCAAUACACUU | | 0.56 $\pm$ 0.03 | 0.50 – 0.60 | 0.36 $\pm$ 0.04 | 0.00 – 0.08 | 8/16 | 0.9999 / 0.9978 | 13/16 | 31 |
| Region 36 | 24368 | 17 | GACUCACUUUCUUCCAC | | 0.68 $\pm$ 0.07 | 0.53 – 0.74 | 0.24 $\pm$ 0.07 | 0.00 – 0.01 | 8/17 | 0.9992 / 0.9877 | 16/17 | 47 |
| Region 37 | 25015 | 18 | GUUAGAUAAAUAUUUUAA | | 0.78 $\pm$ 0.04 | 0.70 – 0.83 | 0.15 $\pm$ 0.03 | 0.00 – 0.03 | 6/18 | 0.9999 / 0.9987 | 14/18 | 11 |
| Region 38 | 27312 | 16 | UAAAAAUUUAUCUAAG | ORF6 | 0.76 $\pm$ 0.01 | 0.74 – 0.77 | 0.17 $\pm$ 0.01 | 0.00 – 0.19 | 7/16 | 0.9999 / 0.9977 | 14/16 | 13 |

**Table S7. Accessible regions by single sequence folding (applying LinearSampling on the SARS-CoV-2 reference sequence alone). The accessibility of the corresponding regions in other 15 SARS-CoV-2 genomes are calculated for each sequence separately. Except for the region in the M gene (in bold), all accessible regions on the reference sequence are not accessible on the other sequences, and always result in a wide range of accessibilities. By contrast, LinearTurboFold is able to find regions that are accessible across all 16 SARS-CoV-2 genomes thanks to fact that consensus folding is determined across the homologous sequences (Tab. S5).**

| Start | Length | Sequence | Gene | Accessibility | | |
|---|---|---|---|---|---|---|
| | | | | Reference Sequence | SARS-CoV-2 sequences (15) | |
| | | | | | Average | Range |
| 9555 | 15 | UUUACUCAUUCUUAC | ORF1ab | 0.61 | $0.52 \pm 0.19$ | 0.00 – 0.62 |
| 20147 | 17 | UCAAUUAUUAUAAGAAA | ORF1ab | 0.56 | $0.07 \pm 0.16$ | 0.00 – 0.55 |
| 23705 | 16 | CCCACAAAUUUUACUA | S | 0.71 | $0.55 \pm 0.31$ | 0.01 – 0.90 |
| 23985 | 15 | AUCCAUCAAAACCAA | S | 0.62 | $0.59 \pm 0.15$ | 0.05 – 0.72 |
| 25700 | 20 | CCCCUUUUCUCUAUCUUUAU | S | 0.97 | $0.17 \pm 0.26$ | 0.00 – 0.98 |
| **27129** | **18** | **AACUAUAAAUUAAACACA** | **M** | **0.77** | $\mathbf{0.76 \pm 0.04}$ | **0.62 – 0.78** |
| 28433 | 15 | ACCGCUCUCACUCAA | N | 0.55 | $0.27 \pm 0.27$ | 0.00 – 0.70 |
| 28691 | 17 | AAUACACCAAAAGAUCA | N | 0.54 | $0.45 \pm 0.22$ | 0.01 – 0.67 |
| 29074 | 16 | AUACAAUGUAACACAA | N | 0.83 | $0.56 \pm 0.40$ | 0.01 – 0.83 |

**SI Dataset S1 (sars-cov-2_and_sars-related_25_genomes_msa_structures.txt)**
Complete results from LinearTurboFold for 25 SARS-CoV-2 and SARS-related genomes.
